# Integrative CRISPR Screens and RNA-Omics Discover an Essential Role for PUF60-3’ Splice Site Interactions in Cancer Progression

**DOI:** 10.1101/2025.05.01.651692

**Authors:** Alexandra T Tankka, Jaclyn M. Einstein, Catherine J. Zhou, Vivian N. Pham, Yuhan Zhang, Jack T. Naritomi, Grady G. Nguyen, Orel Mizrahi, Mark Perelis, Joseph Sarsam, Frederick E. Tan, Dan S. Kaufman, Corina E. Antal, Gene W. Yeo

## Abstract

RNA-binding proteins (RBPs) are important regulators of post-transcriptional gene expression. Understanding which and how RBPs promote cancer progression is crucial for cancers that lack effective targeted therapies such as triple negative breast cancer (TNBC). Here, we employ both *in vitro* and *in vivo* pooled CRISPR/Cas9 screening to identify 50 RBP candidates that are essential for TNBC cell survival. Integrated eCLIP and RNA-sequencing analysis identify that poly(U)-binding splicing factor 60 (PUF60) drives exon inclusion within proliferation-associated transcripts that, when mis-spliced, induce cell cycle arrest and DNA damage. Furthermore, disrupting PUF60 interactions with 3’ splice sites via a substitution in its RNA-binding domain causes widespread exon skipping, leading to downregulation of proliferation-associated mRNAs and inducing apoptosis in TNBC cells. We demonstrate that loss of PUF60-RNA interactions inhibits TNBC cell proliferation and shrinks tumor xenografts, revealing the molecular mechanism by which PUF60 supports cancer progression.

**Significance:** Our work demonstrates functional *in vivo* screening of RBPs as an effective strategy for identifying unexpected cancer regulators. Here, we reveal a crucial role for PUF60-mediated splicing activity in supporting oncogenic proliferation rates and highlight its potential as a therapeutic target in triple negative breast cancer.

## Introduction

RNA-binding proteins (RBPs) govern post-transcriptional gene regulation, a pivotal mechanism exploited by cancer cells to modulate protein expression levels and sustain oncogenic proliferation^1^. RNA splicing, polyadenylation, stability, subcellular localization, and translation are modulated by RBP-RNA interactions^2^. Dysregulated RNA splicing has emerged as a significant driver of cancer progression, as most tumor types display extensively altered splicing compared to healthy tissues^3^. Aberrant expression of splicing regulatory RBPs generates alternatively spliced (AS) isoforms that affect the function, localization, or stability of specific oncoproteins and tumor-suppressors, influencing apoptosis, angiogenesis, proliferation, and metastasis^4^. While it has been recently appreciated that RBP expression and activity is disrupted in various cancers, advances in experimental and computational methods have identified thousands of novel RBPs whose functions in cancer are poorly or incompletely understood^5–9^. Therefore, large-scale functional discovery and characterization of RBPs in cancer is crucial for deepening our understanding of tumor biology and identifying new therapeutic targets.

Triple-negative breast cancer (TNBC) is the deadliest breast cancer subtype, lacking expression of the estrogen receptor (ER), progesterone receptor (PR), and human epidermal growth factor receptor 2 (HER2)^10^. Therapies that block the effects of estrogen and progesterone are effective in treating hormone receptor-positive breast cancers^11^; however TNBCs lack such targeted therapies due to their absence of hormone receptor expression^12^, emphasizing an urgent need for the discovery of effective targets. Accumulating evidence indicates that TNBCs display significant differences in the expression of mRNAs subject to RBP-dependent post-transcriptional control from those in hormone receptor-positive breast cancers and normal breast tissue^13–15^. Moreover, RBPs involved in mRNA splicing and m6A-mediated RNA decay are selectively required for cell survival in TNBC but not for other cell types^16, 17^. These findings emphasize the importance of systematically identifying RBPs that are essential for TNBC tumor growth yet are non-essential in normal tissue-derived cells.

Here, we performed *in vivo* and *in vitro* RBP-focused CRISPR/Cas9 screens to identify RBPs that are specifically required to sustain the survival and proliferation of TNBC cells. We identify 50 RBPs that are required for the survival of TNBC tumors while being dispensable for normal cellular survival. Proteins associated with the U2 small nuclear ribonucleoprotein (U2 snRNP) complex were enriched among the candidates with the poly(U)-binding splicing factor 60 (PUF60) emerging as a key modulator of TNBC proliferation and survival. We found that inhibition of PUF60 binding at splice sites severely restricts cellular proliferation and induces tumor regression in a TNBC xenograft model. PUF60 drives exon inclusion within a network of proliferation-associated transcripts including those involved in cell-cycle progression, chromatin organization, and the DNA damage response. mRNAs within these proliferation-associated pathways undergo widespread exon skipping and destabilization after disrupting PUF60-dependent splice site activity, resulting in TNBC cell apoptosis. Overall, our studies define the functional and molecular role of PUF60-mediated splicing regulation in post-transcriptional gene regulation of cell cycle and genomic stability pathways in TNBC and suggest that PUF60 inhibition is a promising therapeutic approach.

## Results

### Dual CRISPR-Cas9 screens identify RBPs that are required for triple negative breast cancer survival

To identify essential RNA-binding proteins (RBPs) in triple negative breast cancer (TNBC) tumors, we carried out *in vivo* CRISPR/Cas9 screens in tumor xenografts derived from the triple negative breast cancer cell lines MDA-MB-231, MDA-MB-436, and SUM-149. Additionally, we conducted *in vitro* screens in all 3 TNBC lines and the non-transformed MCF10A cell line (“normal”), which is derived from normal breast epithelial tissue and does not form tumors in mice. This dual-screening approach allowed us to identify RBPs essential for the survival of TNBC tumors while excluding those critical for the survival of normal MCF10A cells.

We constructed a single CRISPR RBP-targeting library for use in both *in vitro* and *in vivo* studies. These RBPs were prioritized from our CRISPR/Cas9 drop-out screens performed in isogenic cells containing TNBC-associated genetic alterations. We screened 1,078 RBPs in vitro using isogenic cell lines containing MYC amplification ^16^, KRAS G13D gain-of-function (GoF) (**Table S1**), or heterozygous BRCA1 loss-of-function (LoF) (**Figure S1A-S1B, Table S2**). By compiling all RBP candidates from each of the three screens, we created a new library of 206 RBPs that are essential for at least one TNBC-associated mutation model. Our final CRISPR-Cas9 lentiviral library contained 5 single-guide RNAs (sgRNAs) for each of the targets, alongside 140 sgRNAs targeting 37 essential genes as positive controls, and 225 non-targeting sgRNAs as negative controls (**Table S3**).

MDA-MB-231, MDA-MB-436, and SUM-149 TNBC cell lines as well as normal breast epithelial MCF10A cells were transduced with our CRISPR library, selected with puromycin, and then the TNBC cell lines were subcutaneously transplanted into athymic nude mice (**Figure 1A**). The CRISPR library was constructed in the lenti-CRISPR V2 backbone ^18^, enabling doxycycline (DOX)-inducible expression of Cas9 post-tumor growth to at least 300mm^3^. After 14 days of DOX induction, equatorial tumor cross-sections were harvested, dissociated, and processed through next generation sequencing to quantify pooled sgRNA distribution using the MAGeCK package ^19^. Additionally, all 4 cell lines underwent *in vitro* screening to exclude RBPs that were essential in normal MCF10As from downstream analysis. As expected, following 14 days of DOX-induction *in vitro* and *in vivo*, we observed a reduction in essential gene-targeting sgRNA read counts compared with non-targeting sgRNA read counts, indicating dropout of sgRNA-targeted cells (**Figure S1C**).

**Figure 1.**
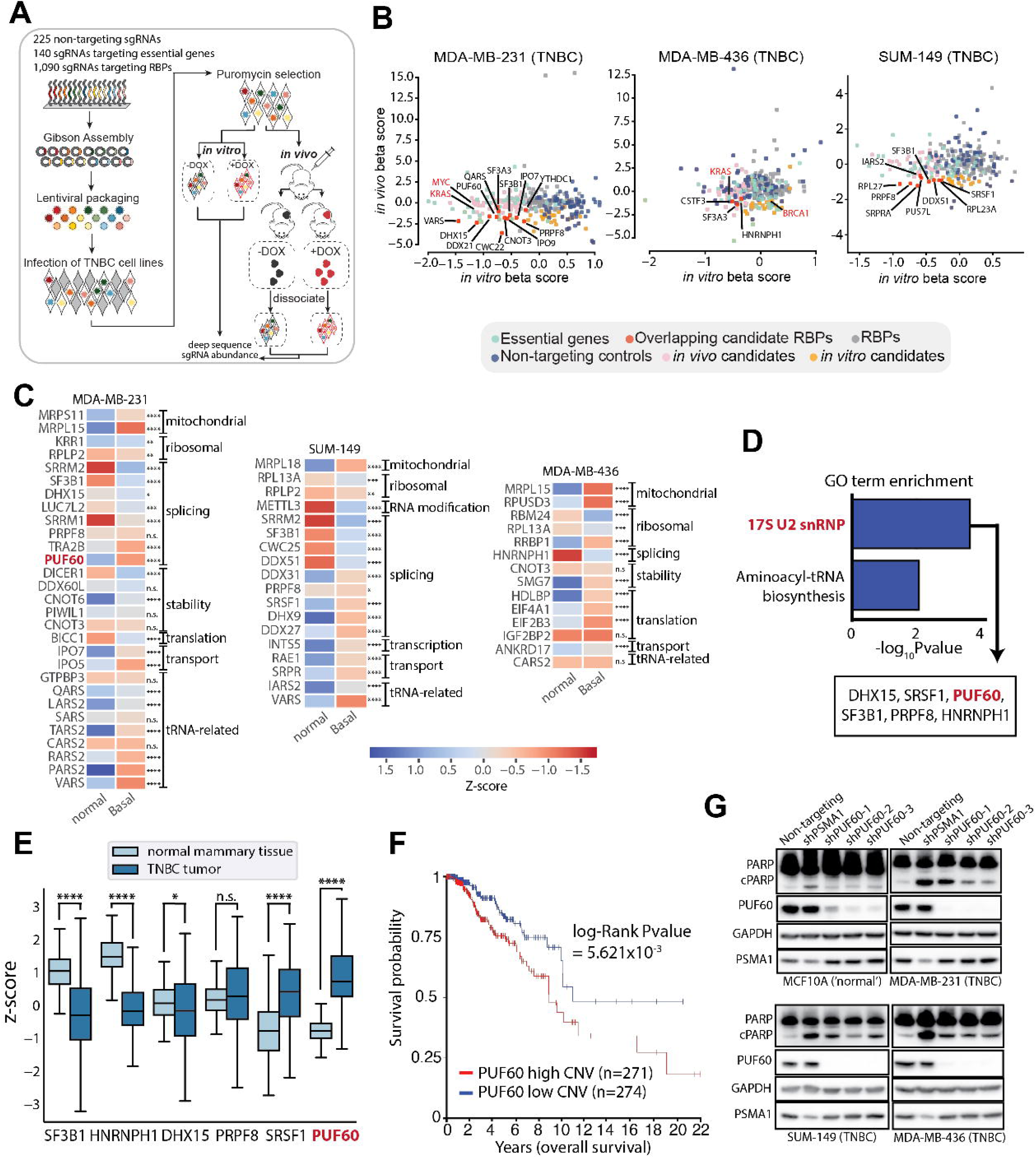
In vivo CRISPR screen identifies U2 snRNP-associated RBPs as an essential vulnerability in TNBC but not non-transformed cells. (A) Schematic of the generation and execution of the RBP-targeted *in vivo* and *in vitro* CRISPR screens in human triple negative breast cancer (TNBC). (B) Comparison of β scores for non-targeting controls (blue), essential genes (green), and RBPs (grey) between *in vitro* and *in vivo* screens. *In vitro* candidates (orange), *in vivo* candidates (pink), and overlapping candidates (red) are indicated. (C) Heat map comparing mRNA expression levels for *in vivo* RBP candidates between normal mammary tissue (GTEx) and Basal-like tumors (TCGA) from the UCSC Xena Browser ^84^. *p<0.05, **p<0.01, ***p<0.001, ****p<0.0001, Student’s t test with Benjamini-Hochberg procedure for multiple comparisons. (D) Gene Ontology (GO) enrichment of overlapping RBP candidates between in vitro and in vivo screens. (E) Boxplots comparing normal mammary tissue (GTEx) and Basal-like tumor (TCGA) mRNA expression levels of overlapping RBP candidates belonging to the 17S U2 snRNP ontology term. *p<0.05 and ****p<0.0001, Student’s t test with Benjamini-Hochberg procedure for multiple comparisons. (F) Kaplan-Meier survival analysis of breast cancer survival probability comparing tumors with lowest quartile (blue, n=274) PUF60 copy number variation (CNV) and highest quartile (red, n=271) PUF60 CNV. LogRank significance. (G) Western blot analysis of cell lysates from non-targeting control (NTC), shPSMA1, and shPUF60 TNBC and normal mammary epithelial cell lines showing cleaved PARP protein expression.

Computational analysis revealed that 69 RBPs (Wald p < 0.05) were significantly depleted in DOX-induced TNBC tumors versus uninduced control tumors. Of these, 50 RBP candidates were specifically depleted in DOX-induced TNBC tumor cells compared with DOX-induced MCF10A cells, indicating unique roles in TNBC survival (**Figure 1B, Table S4**). This cohort included some RBPs with previously reported roles in TNBC-associated RNA metabolism such as IGF1BP2^20^, SF3B1^21^, METTL3^22^, and TRA2B^23^. We also identified 40 RBPs that have not yet been explored in TNBC, alongside several candidates with unknown function in any oncogenic context including CARS2, RPUSD3, SRPR, INTS5, PARS2, and GTPBP3. Next, we analyzed public data from The Cancer Genome Atlas (TCGA) pan-cancer clinical data resource^24^ and The Genotype-Tissue Expression (GTEx) comprehensive tissue resource^25^ and observed that 29 out of 50 RBP candidates are highly expressed in basal-like tumors (**Figure 1C**), a subtype associated with triple-negative characteristics^26^. To define a high-confidence cohort of RBPs, we identified a core set of 15 TNBC-essential candidates significantly depleted in both *in vitro* and *in vivo* DOX-induced TNBC cells. GO ontology analysis highlighted enrichment of U2 small nuclear ribonucleoprotein complex (U2 snRNP) components (PUF60, DHX15, SRSF1, PRPF8, HNRNPH1, SF3B1) among these high-confidence candidates (**Figure 1D and Figure S1D-E**). Consistent with literature indicating high mutation frequencies or aberrant expression of U2snRNP components across cancers ^27–29^, our results suggest that TNBC tumors are particularly dependent on RBPs that either recruit U2snRNP to 3’ splice sites (3’SS) or integrate directly into the complex to influence branchpoint usage.

### Depleting PUF60 induces apoptosis of TNBC cells

Among the U2 snRNP-associated RBPs, transcript levels of the poly(U)-binding splicing factor PUF60 were the most significantly upregulated in basal-like tumors compared to normal breast tissue (**Figure 1E**). PUF60 functions as a splicing factor by directly binding polypyrimidine tracts (PPTs) to promote U2 snRNP association with 3’SSs in pre-mRNA of weak splice sites^30, 31^. It is located within the 8q24 chromosomal region, alongside the MYC oncogene, which is one of the most frequently amplified regions in TNBC. This region contains risk loci for multiple epithelial malignancies including breast cancer^32–34^. Thus, previous work has reported copy number gain or elevated expression of PUF60 in various tumor types, along with roles in cancer progression^35–39^. However, whether these cancer-promoting functions necessitate MYC amplifications remains insufficiently explored. Although PUF60 is a nucleic-acid binding protein involved in nuclear splicing and transcriptional processes^30, 40^, only one prior study has explored PUF60-mediated splicing activity in cancer ^38^. The majority of studies propose that its cancer-related functions primarily involve cytoplasmic interactions ^37, 39^ or RNA- and DNA-independent activity ^35^. Furthermore, the transcriptome-wide direct binding and splicing targets of PUF60 remain undefined in any cell or tissue type, emphasizing the need to investigate its direct, splicing-regulatory targets in TNBC.

To explore the clinical relevance of PUF60, we cross-referenced the TCGA Breast Cancer (BRCA) dataset and observed significantly improved overall survival of patients with breast cancer tumors in the lowest quartile of PUF60 copy number variation (CNV) compared to breast cancer patients in the highest quartile (**Figure 1F**). Consistently, we confirmed significant upregulation of PUF60 transcript levels in TNBC cell lines-MDA-MB-231, MDA-MB-436, and SUM-149 compared to normal MCF10A cells (**Figure S1F**).

Subsequently, using three independent short hairpin RNA (shRNA)-mediated knockdowns of PUF60 (shPUF60-1, shPUF60-2, and shPUF60-3), we demonstrated that loss of PUF60 expression induced apoptosis as evidenced by a selective increase of PARP-1 cleavage only in TNBC cell lines while normal epithelial breast cells showed no change (**Figure 1G**). In contrast, knockdown of the essential RBP, PSMA1 (shPSMA1), resulted in PARP-1 cleavage across all cells tested. These results highlight that PUF60 is required for TNBC survival but dispensable for the survival of normal epithelial breast MCF10A cells.

### PUF60 targets are enriched for genes regulating cell cycle progression, the DNA damage response, and chromatin remodeling

To identify PUF60 splicing regulatory targets in TNBC, we performed enhanced crosslinking and immunoprecipitation (eCLIP) ^41^ in MDA-MB-231, MDA-MB-436, SUM-149, and MCF10A cells, in biological duplicate (**Figure S2A**). Consistent with previous in vitro splicing assays indicating that PUF60 facilitates the formation of U2snRNP complexes ^30^, we observed a large fraction of reproducible enriched PUF60 binding windows immediately adjacent to the 3’SS (3’SS ADJ) (**Figure 2A and S2B**). Metagene analysis of eCLIP data confirmed enrichment at the 3’SS across all cell lines (**Figure 2B**). There is no consensus motif known to influence PUF60’s selection of PPTs and, while PPTs are generally known to be uracil-rich, there is considerable variability in the specific sequence of a given PPT ^42^. Notably, motif analysis on the reproducible enriched windows from each cell line revealed a highly significant alternating UC residue from the eCLIP signal, suggesting a sequence preference for PUF60 at PPTs containing a UCUCUCUC motif (**Figure 2C**).

**Figure 2.**
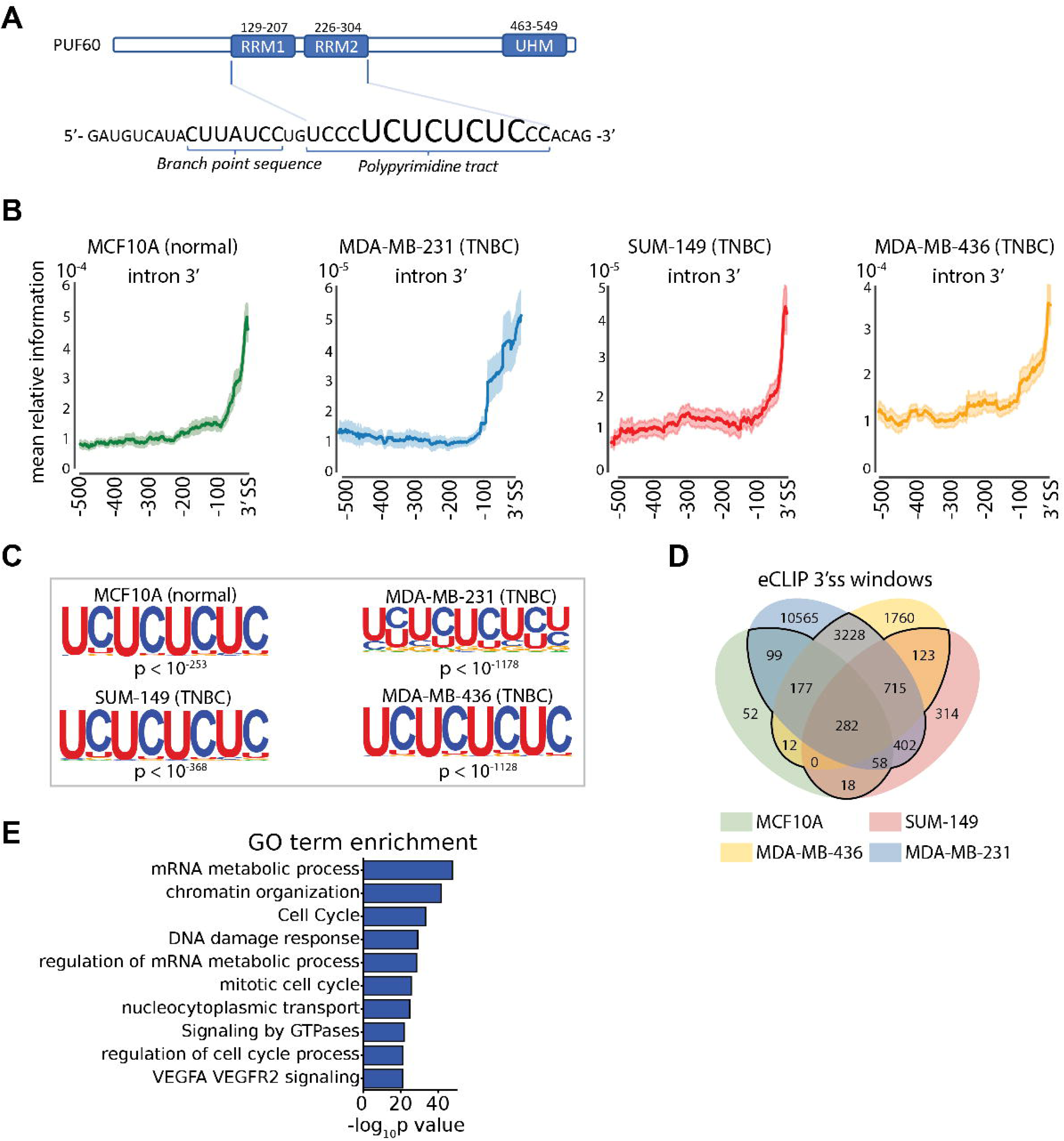
eCLIP reveals enriched PUF60 binding at 3’ splice sites within cell-cycle, chromatin organization, and DNA damage-associated transcripts. (A) Domain structures of PUF60 and the pre-mRNA 3’ intron sequence. (B) Metadensity profiles of unfiltered eCLIP peak enrichment at 3’ splice sites (3’SS) ^79^. (C) Motif analysis of eCLIP data. Representative of two independent replicates (D) Four-way Venn diagram of PUF60 3’SS target exons among TNBC and normal epithelial breast cell lines. 3’SS targets bound by PUF60 in at least two cell lines are outlined. (E) Gene Ontology (GO) enrichment of genes containing 3’SSs bound by PUF60 in at least two cell lines as outlined in (E).

To investigate PUF60’s role in TNBC survival and distinguish specific pathways that contribute to apoptosis upon PUF60 depletion, we compared high-confidence 3’SS binding sites among TNBC and normal cell lines. Given that RBP binding activity is generally regulated by RNA sequences and secondary structures, which are conserved between cell types and states, we found a significant overlap of 3’SS binding sites among each cell line (**Figure S2C**). We defined a common set of 4,991 3’SS eCLIP binding sites (black outline) occurring in at least two out of four cell lines (**Figure 2D, Table S5**) and observed enrichment for genes that regulate cell cycle progression, the DNA damage response, and chromatin organization (**Figure 2E**). Unlike in healthy tissues, cancer cells undergo continuous rounds of division, increasing their reliance on cell cycle-dependent processes including the DNA damage response and chromatin remodeling ^43^. Our findings indicate that PUF60 binds to the 3’SS of transcripts associated with these cancer-promoting processes, implicating a potential role for PUF60 in sustaining high rates of oncogenic proliferation.

### Loss of PUF60 downregulates cell cycle and DNA damage-associated transcripts

We next investigated the impact of PUF60 knockdown on alterative splicing (AS) by performing RNA sequencing (RNA-seq) on PUF60-depleted TNBC and normal cell lines in biological duplicate (**Figure S3A-B**). Globally, we detected a large proportion of exon skipping events compared to all other AS events (**Figure 3A and S3C**) which is consistent with prior knockdown RNA-seq studies ^44^. Among PUF60 3’SS targets, we also observed decreased exon inclusion upon PUF60 knockdown, confirming that PUF60 binding upstream of an exon promotes inclusion of that exon (**Figure 3B**).

**Figure 3.**
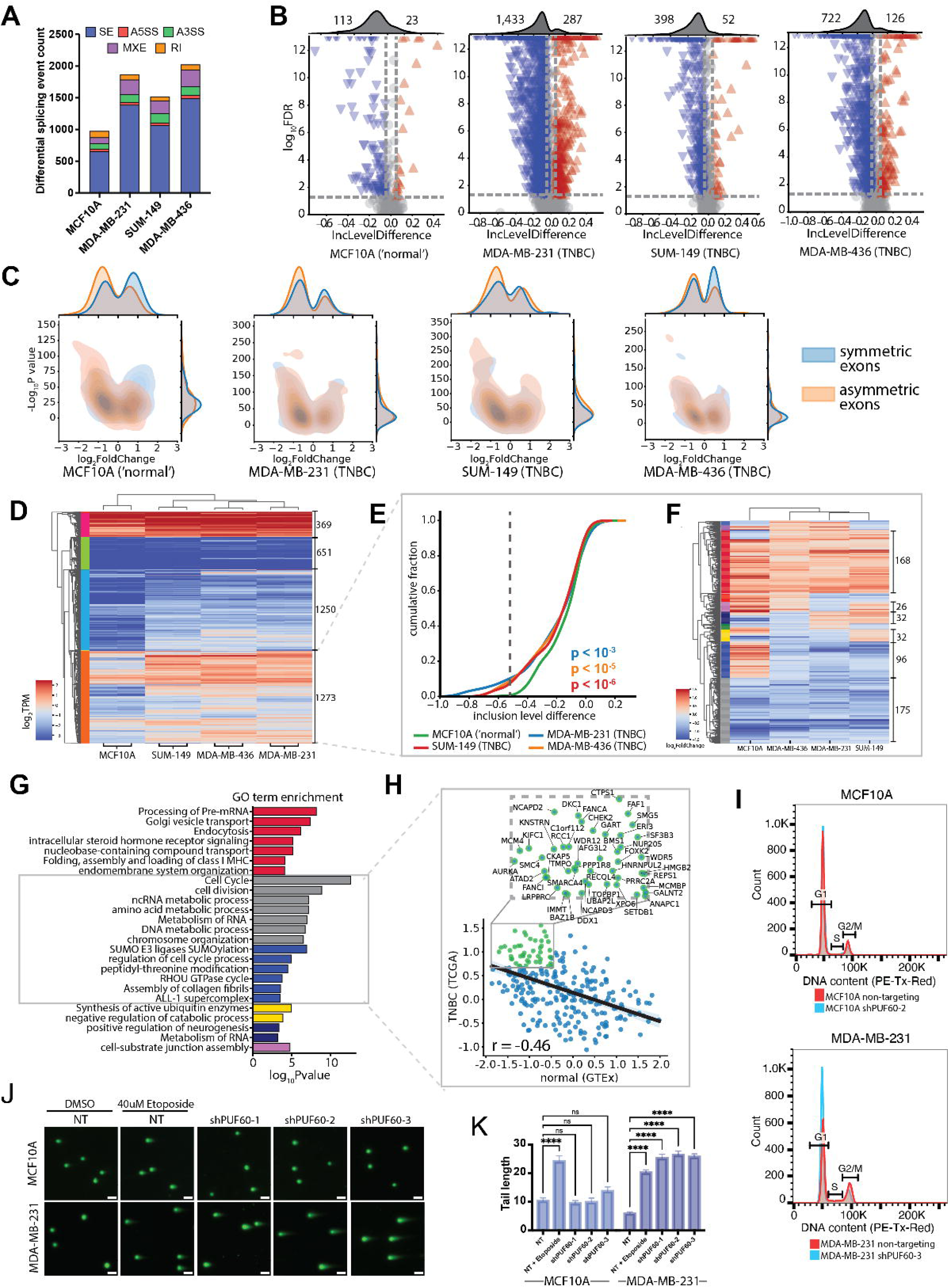
PUF60 regulates the splicing of cellular proliferation transcripts that are highly expressed in TNBC. (A) Stacked bar plot of differentially spliced events after shRNA-mediated knockdown of PUF60 as identified by rMATS analysis. Differential splicing events have an inclusion levels difference > 0.05 or < −0.05 and a false discovery rate (FDR) < 0.01 as calculated by the rMATS log likelihood ratio. (B) Volcano plots describing the included (red, inclusion level difference > 0) and excluded (blue, inclusion level difference < 0) PUF60 KD-sensitive exon splicing events bound by PUF60 at the 3’ splice site (3’SS). Black lines with grey shading (top) summarize the distribution of PUF60 splicing regulatory events by exon inclusion and exclusion. (C) Jointplots illustrating the expression of genes containing KD-sensitive exons bound by PUF60 as identified by DESeq2 analysis. Genes either contain KD-sensitive symmetric exons (blue) or KD-symmetric asymmetric exons (orange). (D) Hierarchical cluster map illustrating exon expression levels of overlapping PUF60 3’SS targets outlined in (Fig. 2F). Exon expression levels calculated by featurecounts of RNA-seq data from unperturbed cells. Gene clusters are depicted in the dendrogram. (E) Cumulative distribution of inclusion level differences from exons within the red cluster from (**D**). Kolmogorov-Smirnov test. (F) Hierarchical cluster map describing differential expression of genes containing exons within the red cluster from (**D**) as determined by DESeq2 analysis. Differentially expressed genes following PUF60 knockdown have a DESeq2-determined adjusted P value < 0.05. (G) Gene Ontology (GO) enrichment of genes in the dendrogram clusters from (**F**). Red, grey, blue, yellow, navy, and purple bar colors correspond to dendrogram cluster colors. (H) Regplot comparing mRNA expression level Z scores of genes within grey and blue clusters from (**F**) between basal-like breast cancer (TCGA data portal) ^24^ and normal breast tissues (GTEx comprehensive tissue resource) ^25^. Inset highlights the most upregulated PUF60 targets in TNBC tumors exhibiting downregulated expression in normal tissues. Pearson’s correlation test. (I) Overlayed histograms of PI staining by flow cytometry to assess DNA content of MCF10As (top) and MDA-MB-231s (bottom) expressing non-targeting control shRNA (red) and PUF60-targeting shRNA (blue) (J) Representative comet assay images at 20x magnification from MCF10As and MDA-MB-231s expressing non-targeting control shRNA and PUF60-targeting shRNAs. Scale bar = 100 μm (K) Bar plots illustrating quantification of tail length. ****p<0.0001, two-way ANOVA with Dunnett’s test for multiple comparisons.

Next, we verified the functional consequences of PUF60 depletion and found that, following PUF60 knockdown, the majority of PUF60 targets containing exon skipping events were downregulated in expression (**Figure S3D**). We hypothesized that the decrease in PUF60 target transcript expression was caused by asymmetric exons (exon nucleotide length not divisible by 3), or exons that disrupt the downstream reading frame when removed, potentially triggering RNA surveillance pathways such as nonsense-mediated decay (NMD). Indeed, across all four cell lines, over 50% of differentially expressed PUF60 regulatory targets were asymmetric exons. Moreover, a higher proportion of transcripts with asymmetric PUF60 regulatory targets were downregulated following knockdown compared to transcripts with symmetric targets (**Figure 3C**). Our findings suggest that knockdown of PUF60 in both TNBC and normal cells destabilizes transcripts because they contain skipping events within asymmetrical exons.

To identify PUF60 targets with cancer-specific survival roles, we analyzed the expression level of exons containing a 3’SS peak in at least two cell lines from unperturbed MCF10A and TNBC cells (black outline in **Figure 2E**). Using hierarchical clustering of normalized transcripts per million reads (TPMs), we observed that the largest cluster (red, 1273 genes) consisted of PUF60-bound exons with higher expression levels in TNBC under basal conditions (**Figure 3D, Table S6**). Genes with exons within this cluster exhibited a significant increase in the magnitude of PUF60 KD-sensitive exon skipping events in TNBC compared to normal (**Figure S3E and 3E**), indicating that PUF60 depletion results in more robust exon skipping in transcripts that are highly expressed in TNBC. We then performed hierarchical clustering of the differentially expressed PUF60 targets within the orange cluster (**Figure 3D**) to identify pathways uniquely contributing to apoptosis after PUF60 depletion (**Figure 3F, Table S7**). Two clusters (blue and grey) showed downregulation of PUF60 targets across all TNBC cell lines. These clusters were enriched for genes regulating cell cycle, chromosome organization, and collagen fibril assembly, all of which are dynamically regulated during cellular proliferation (**Figure 3G**). Our findings indicate that depletion of PUF60 downregulates genes that encode and regulate cellular proliferation through exon skipping events within overexpressed genes in TNBC. Importantly, only TNBC cells were highly sensitive to downregulation of these pathways, possibly due to their increased dependence on cell cycle control to maintain continuous proliferation.

To explore the clinical relevance of PUF60 targets that showed downregulation upon knockdown (blue and grey clusters (**Figure 3H**), we integrated RNA-seq data from TCGA pan-cancer clinical data resource ^24^ and GTEx comprehensive tissue resource ^25^. Hierarchical clustering of Z scores revealed that most genes within this cohort were overexpressed in TNBC tumors compared with normal breast tissue (139 genes) (**Figure S3F**). Additionally, we found a negative correlation (R^2^ = −0.46) between gene expression levels in TNBC tumors and normal breast tissue, indicating that highly expressed PUF60 targets in tumor samples are generally lowly expressed in normal breast tissue (**Figure 3H**). The most upregulated PUF60 targets in TNBC tumors, which exhibited downregulated expression in normal tissues, included MCM4^45^, CHEK2^46^, NCAPD2^47^, RCC1^48^, and SETDB1^49^, all of which are implicated in the DNA damage response and cell cycle progression in various cancers.

Given PUF60’s role in splicing regulation of cell cycle transcripts, we used propidium iodide (PI) staining to determine the cell cycle status of TNBC and normal cells after PUF60 knockdown. Notably, PUF60 depletion in MDA-MB-231 cells led to a significant increase in the proportion of cells in the G1 phase, and a corresponding decrease in the G2/M phase. MCF10As did not display robust changes in G1 or G2/M phase distribution following PUF60 depletion (**Figure 3I and S3G**). Although we found that depletion of PUF60 decreases cell cycle-associated transcript expression in both TNBC and normal cells, only TNBC cells–characterized by unchecked proliferation–were highly reliant on PUF60 for proper cell cycle progression. To further characterize the importance of PUF60’s regulatory targets associated with DNA damage, we performed a comet assay to quantify DNA damage resulting from PUF60 depletion in both TNBC and normal cells. Indicators of DNA damage, including the percentage of DNA in the tail, tail moment, and tail length were all elevated in TNBC cells following PUF60 knockdown while these parameters were unchanged in normal cells (**Figure 3J-K and S3H-I**). Our findings suggest that TNBC cells are much more susceptible to DNA damage after PUF60 knockdown, contributing to the essential role of PUF60 for TNBC cell survival. In summary, PUF60 binds to and regulates the splicing of key transcripts to prevent DNA damage and cell cycle arrest–processes which are typically evaded in highly proliferative cancer cells.

### Depleting PUF60 induces TNBC tumor regression and triggers apoptosis *in vivo*

To assess whether PUF60 depletion negatively affects tumor growth in vivo, we established stable MDA-MB-231 cells transduced with 3 distinct DOX-inducible PUF60 shRNAs. DOX treatment effectively reduced PUF60 protein level for all three shRNAs (**Figure S4A**). Following subcutaneous engraftment of MDA-MB-231 cells expressing 2 of our validated shRNAs and subsequent DOX administration (**Figure 4A)**, we observed an 86% reduction in tumor burden compared to vehicle-treated controls (**Figure 4B, 4C, and S4B**). Protein lysates from these tumors confirmed sustained PUF60 depletion throughout the treatment period (**Figure 4D**). To better mimic the clinical scenario observed in patients, we next assessed the therapeutic potential of PUF60 inhibition in inducing TNBC tumor regression by initiating DOX treatment only after tumors reached a larger size (average volume of 132.8 mm^3^). Following 29 days of DOX treatment, tumor volume decreased by 53% (**Figure 4E, 4F, Table S8**), demonstrating that PUF60 not only inhibits tumor growth but also induces tumor regression. Fluorescent *in vivo* imaging of the RFP-tagged tumors further corroborated these findings, showing loss of RFP signal by day 47 of DOX treatment (**Figure S4C-D**). Protein lysates from these tumors (**Figure S4E**) confirmed PUF60 depletion (**Figure S4F**). Additionally, we detected a significant increase in cleaved caspase-3 (CC3) positive cells in tumor sections from DOX-treated mice, indicating that PUF60-depletion leads to tumor regression via induction of apoptosis in TNBC tumors (**Figure 4G-H**). Our findings collectively indicate that PUF60 is required to sustain growth and survival in TNBC tumors.

**Figure 4.**
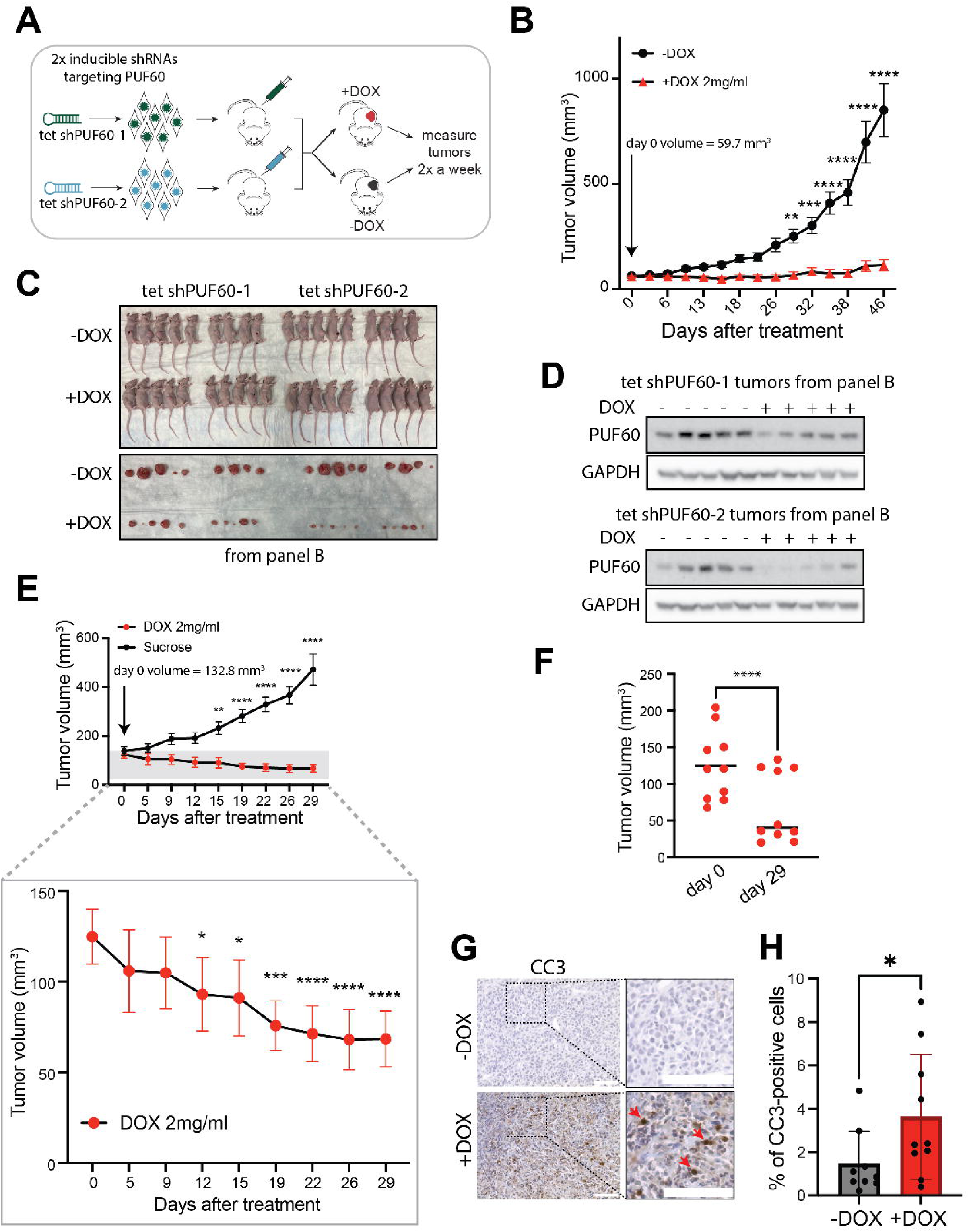
Depletion of PUF60 in TNBC cells significantly inhibits tumor growth. (A) Schematic depicting the generation of doxycycline (DOX)-inducible PUF60 knockdown xenograft growth curves. (B) Average tumor volume over 46 days of DOX-induced (red), PUF60-depleted xenografted mice compared with vehicle controls (black). Initial (day 0) average tumor volume is 59.74 mm^3^. **p<0.01, ***p<0.001, ****p<0.0001, two-way ANOVA with Bonferroni’s test for multiple comparisons. Bars are mean ± SEM; -DOX, n = 18 mice; +DOX, n = 20 mice. (C) Images of mice and excised tumors 46 days following DOX induction compared with vehicle controls. (D) Western blot analysis of cell lysates from final tumors from (**B**) showing PUF60 protein expression in DOX-induced compared to vehicle control tumors. (E) Average tumor volume of DOX-induced tumors with initial (day 0) average volume of 132.78 mm^3^. Statistical comparisons between +DOX and -DOX tumor volumes for each time point. Inset highlights DOX-treated tumor volume with reduced y axis range. Statistical comparisons between each time point and initial (day 0) tumor volume, *p<0.05, **p<0.01, ***p<0.001, ****p<0.0001, two-way ANOVA with Bonferroni’s test for multiple comparisons. Bars are mean ± SEM; -DOX, n = 12 mice; +DOX, n = 10 mice. (F) Tumor volumes of DOX-induced tumors at initial (day 0) compared with 29 days following treatment. ****p<0.0001, paired Student’s t test. (G) Representative 40x images of immunohistochemical cleaved caspase 3 (CC3) staining of tumor sections from mice treated with DOX or vehicle for 45 days. Dashed boxes are magnified in the insets. Red arrows indicate CC3 staining. All scale bars, 90 μM. (H) Bar plot summarizing the percentage of CC3-positive cells. Each point represents the mean percentage of CC3-positive cells for an individual tumor. *p<0.05, one-sided Mann Whitney test. Bars are mean ±SD; -DOX, n = 9 tumors; +DOX, n = 10 tumors

### A residue within PUF60’s RRM1 domain is required for global recognition of 3’ splice site targets

While previous studies have proposed that PUF60’s cancer-related functions primarily involve cytoplasmic protein-protein interactions^37, 39^ or nucleic acid-independent activities^35^, we observed predominantly nuclear signal in MDA-MB-231 cells immunostained for PUF60 (**Figure S5A**). To evaluate the role of PUF60’s RNA-binding activity in TNBC, we investigated if disrupting its ability to bind 3’SSs would replicate the effects observed in our general knockdown studies. We constructed TNBC cells with an L140P mutation (L140P) within the RNA recognition motif 1 (RRM1) domain of PUF60, which Kralovicova et. al previously characterized to impair exon inclusion at PUF60-activated splice sites by reducing the ability of PUF60 to bind it’s U-rich substrates^50^. To investigate the RNAs bound by this mutant, we transduced MDA-MB-231s expressing an endogenous PUF60 shRNA (tet-shPUF60-1) with wild-type (WT-PUF60) or L140P (L140P-PUF60) codon-optimized and V5-tagged PUF60 (**Figure 5A and S5B-C**). We then performed eCLIP on L140P-PUF60 and WT-PUF60 MDA-MB-231s using a V5 antibody. Our *de novo* motif analysis of WT-PUF60 reproducible enriched eCLIP windows identified the same significantly enriched UCUCUCUC motif (p <10^-1^^31^) detected in our endogenous PUF60 eCLIPs. In contrast, enrichment for this motif was absent in the L140P-PUF60 eCLIP signal. Furthermore, we observed significant enrichment for uracil-rich sequences in WT-PUF60 eCLIP windows, aligning with PUF60’s established role in directly binding PPTs, whereas uracil enrichment was lacking in L140P-PUF60 eCLIP windows (**Figure 5B**). Despite distinct motif patterns observed between L140P- and WT-PUF60 eCLIP signals, we found that canonical donor (AG’**GU**RAGU) and acceptor (Y**AG**) splice sites were enriched among L140P-PUF60 motifs (**Figure 5B**). While our analysis indicates that L140P-PUF60 is not interacting with PPTs, our findings suggest that the RRM1 mutant may be recruited to splice sites through protein-protein interactions with U2snRNP components, facilitated by L140P-PUF60’s intact U2AF homology motif (UHM) domain.

**Figure 5.**
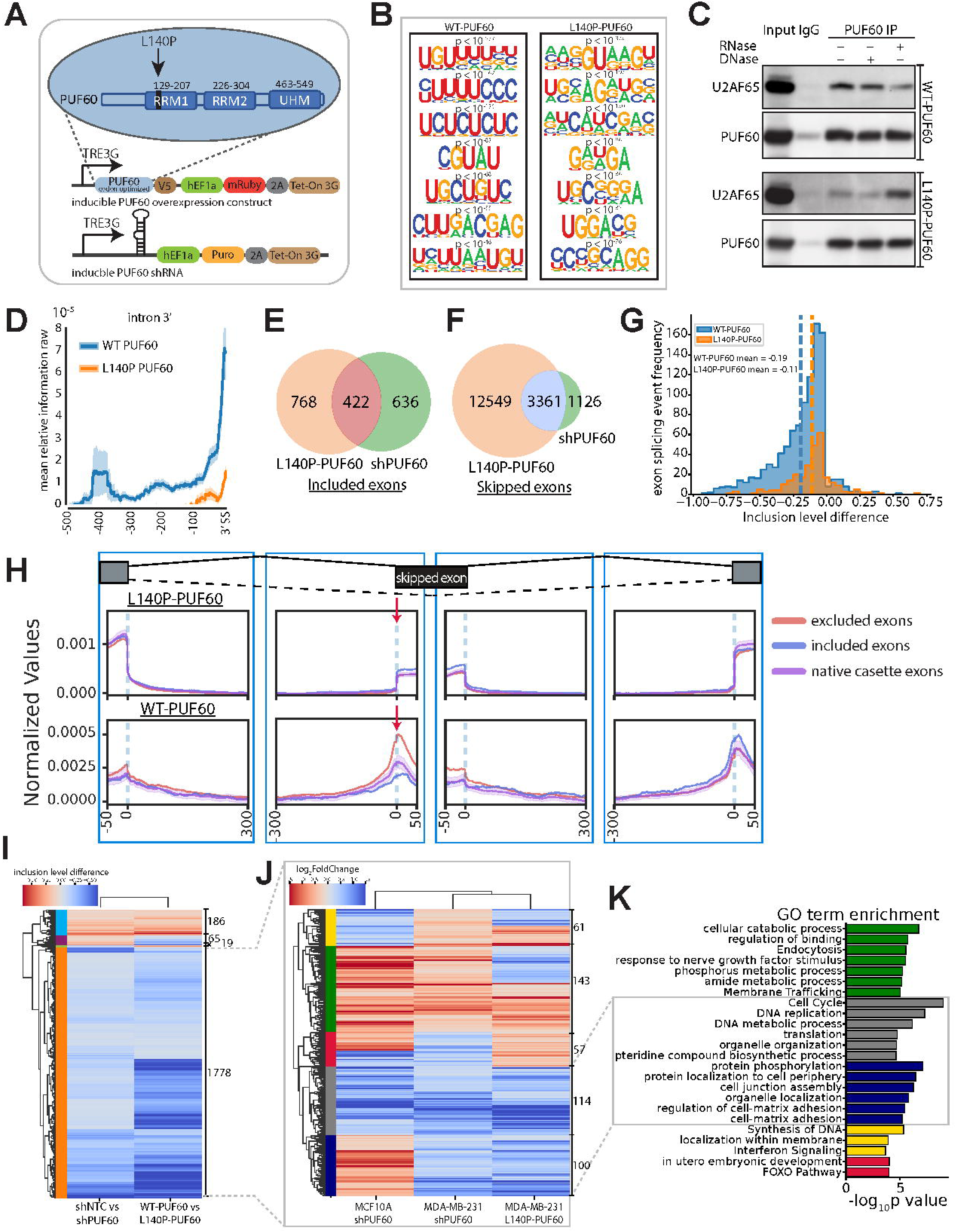
A substitution within PUF60’s RRM1 domain downregulates cellular proliferation pathways. (A) Schematic describing generation of stable L140P-PUF60 and WT-PUF60 cells. An inducible, codon-optimized open reading frame (ORF) encoding either L140P or wild-type (WT) PUF60 is stably expressed along with an inducible shRNA specific to endogenous PUF60 (bottom) in the MDA-MB-231 TNBC line. (B) Top 7 most significantly enriched RNA binding motifs as identified by HOMER analysis of L140P-PUF60 and WT-PUF60 eclip signal. Corresponding P values calculated by HOMER default settings. Representative of two independent replicates. (C) Western blot of PUF60 immunoprecipitation in L140P-PUF60 and WT-PUF60 MDA-MB-231s, showing co-immunoprecipitation of U2AF65 with and without RNAse A or DNAse treatment. (D) Metadensity profiles of unfiltered L140P-PUF60 and WT-PUF60 eCLIP peak enrichment at 3’ splice sites (3’SS) ^79^. (E) Venn diagram comparisons of differential exon inclusion events after L140P substitution (orange) and PUF60 knockdown (green). Differential exon inclusion events have an inclusion levels difference > 0.05 and a false discovery rate (FDR) < 0.01 as calculated by the rMATS log likelihood ratio. (F) Venn diagram comparisons of differential exon skipping events after L140P substitution (orange) and PUF60 knockdown (green). Differential exon inclusion events have an inclusion levels difference < −0.05 and a false discovery rate (FDR) < 0.01 as calculated by the rMATS log likelihood ratio. (G) Histogram summarizing the frequency of WT-PUF60 (blue) and L140P-PUF60 (orange) 3’SS eCLIP peaks proximal to exon splicing events. Average inclusion level difference of splicing events near WT-PUF60 and L140P-PUF60 peaks is shown. (H) RBP-maps output for L140P-PUF60 (top) and WT-PUF60 (bottom) RNA binding sites proximal to cassette exons and differentially excluded exons following general PUF60 knockdown. Purple shading around ‘native cassette exon’ line indicates 95% confidence interval determined by RBP-maps. (I) Hierarchical cluster map of inclusion level differences from exons containing proximal endogenous PUF60 eCLIP peaks. Differential exon splicing events determined by rMATS are following general PUF60 knockdown and L140P-PUF60. Gene clusters are depicted in the dendrogram. (J) Hierarchical cluster map illustrating differential expression of genes containing exons depicted in the yellow cluster in (**I**). Genes have a DESeq2-determined adjusted P value < 0.05 (K) Gene Ontology (GO) enrichment of genes with downregulated expression depicted in blue and grey clusters in (**J**)

In line with this hypothesis, co-immunoprecipitation (Co-IP) assays revealed that L140P-PUF60 directly interacts with the U2-associated splicing factor, U2AF65, however with lower affinity compared to WT-PUF60 (**Figure 5C**)^31^. Interestingly, we observed a reduction only in WT-PUF60-U2AF65 interactions upon RNase A treatment, aligning with the rationale that PUF60-U2AF65 interactions are, in part, RNA-mediated. Our observations indicate that while L140P-PUF60 can bind U2snRNP proteins localized at splice sites, its ability to engage in RNA-mediated interactions may be reduced due to its decreased binding capability at U-rich RNA substrates. Next, we assessed the relative enrichment of L140P- and WT-PUF60 at 3’SSs and found that WT-PUF60 exhibited approximately 4-fold higher reproducible eCLIP signal in this region compared to L140P-PUF60 (**Figure 5D**). Furthermore, 66% of genes bound by WT-PUF60 exhibited a loss of 3’SS signal after RRM1 mutation, (**Figure S5D-E**) confirming that L140P substantially reduces PUF60’s RNA binding affinity and presence at 3’SSs.

We further investigated the impact of L140P on AS by performing RNA-seq on L140P- and WT-PUF60 MDA-MB-231s (**Figure 5A and S5F**). We initially verified that the L140P substitution excludes the PUF60-activated UBE2F exon 5, a splicing event used to classify L140P as a mutation that suppresses PUF60’s splicing activity ^50^. In line with this, we observed that the presence of the UBE2F exon 5 was nearly eliminated (rMATS FDR = 9.67×10^-4^) following L140P mutation (**Figure S5G-I**). We next assessed whether differential PUF60 knockdown and L140P mutation exon skipping events overlapped, and found that ∼75% of differential exon skipping events observed after general PUF60 KD were also detected upon L140P mutation (**Figure 5E-F**). The remaining 25% of exon skipping events not observed after L140P mutation may stem from indirect effects associated with the non-RNA binding functions of PUF60.

To determine if L140P-PUF60 binding impacts exon usage, we assessed the frequency of WT- and L140P-PUF60 3’SS eCLIP peaks proximal to exon splicing events. Our analysis revealed that WT-PUF60 bound near substantially more exon skipping events, which were also greater in magnitude (WT-PUF60 average inclusion level difference = −0.19, L140P-PUF60 average inclusion level difference = −0.11), than skipping events near L140P-PUF60 peaks (**Figure 5G**). Our findings indicate that L140P-PUF60 does not direct AS of its targets. We further investigated the position-specific impact of L140P-PUF60 binding on neighboring exon usage by constructing a “splicing map” outlining eCLIP signal near skipped exons compared to all native cassette exons ^51^. Our analysis revealed that WT-PUF60 3’SS binding showed global enrichment near skipped exons compared to native cassette exons (**Figure 5H**). Such an enrichment pattern was not observed for L140P-PUF60, further supporting that the L140P substitution disrupts 3’SS-dependent splicing regulation of PUF60’s targets.

### Disruption of PUF60’s 3’ splice site-dependent splicing activity downregulates cellular proliferation pathways

To assess whether there are consistent differential splicing patterns in TNBC cells following PUF60 knockdown and L140P mutation, we examined splicing events involving exons bound by endogenous PUF60 at their 3’SS. Utilizing hierarchical clustering of inclusion levels, we identified that 86% of PUF60’s target exons clustered together (gold) showing exon skipping after both endogenous PUF60 KD and L140P mutation (conserved PUF60 targets), indicating substantial similarity in splicing outcomes between both conditions (**Figure 5I, Table S9**). We next verified that L140P mutation also caused downregulation of cellular proliferation pathways by conducting hierarchical clustering of differentially expressed and conserved PUF60 targets (**Figure 5J, Table S10**). Our analysis identified two clusters (grey and blue) in which conserved PUF60 targets were downregulated both after L140P mutation and general PUF60 knockdown. Unlike clusters containing upregulated genes after PUF60 mutation or knockdown, the grey cluster was enriched for genes associated with cell cycle and DNA metabolism (**Figure 5K**). These findings are consistent with our endogenous PUF60 eCLIP and knockdown RNA-seq data, which identify PUF60 as a regulator of transcripts associated with genomic integrity and cell cycle progression (**Figure 3G**). Notably, the blue cluster showed PUF60 targets that were downregulated in TNBC but upregulated in normal cells, and these targets were associated with cellular matrix adhesion and junction assembly. The ability of cancer cells to communicate with the extracellular matrix (ECM) through cellular adhesions is crucial for intracellular survival and proliferation signaling ^52^, which also supports with our understanding of PUF60’s role in cancer cell division (**Figure 5K**). We conclude that disrupting PUF60’s ability to bind the 3’SS replicated the molecular effects observed in our general knockdown studies, suggesting that PUF60-mediated splicing activity is vital for the regulation of transcripts encoding cellular proliferation regulators.

### Silencing of PUF60 splicing activity by L140P mutation suppresses tumor growth, *in vivo*

To determine the functional implications of PUF60-mediated 3’SS recognition, we measured proliferation rates upon DOX induction of L140P-PUF60 and WT-PUF60 in PUF60-depleted MDA-MB-231 cells (**Figure 5A**). After 48 hours of treatment, we observed that TNBC cells expressing L140P-PUF60 ceased proliferation, whereas cells expressing WT-PUF60 continued to proliferate at rates similar to those of the untreated cells (**Figure 6A**). Notably, L140P-PUF60 MDA-MB-231 cells also exhibited increased levels of caspase 3/7-positive cells (**Figure 6B-C**), underscoring the essential role of PUF60-RNA binding in sustaining TNBC survival, *in vitro*. To corroborate this finding *in vivo*, we subcutaneously transplanted L140P-PUF60 and WT-PUF60 MDA-MB-231s into mice (**Figure 6D and S6A**). Upon tumor engraftment and DOX treatment, we observed a 93% reduction in tumor burden from mice engrafted with L140P-PUF60 cells, compared to mice engrafted with WT-PUF60 cells (**Figure 6E-G, Table S11**). Furthermore, we observed a 67% reduction in L140P-PUF60 tumor size after 58 days of treatment compared with tumor volume on the first day of treatment (**Figure 6F and 6H**). Our observations indicate that, in addition to suppressing tumor growth, the PUF60 L140P mutation effectively induces tumor regression. Fluorescent *in vivo* imaging of RFP-tagged tumors treated with DOX revealed a loss of RFP signal by day 47 of treatment, despite the presence of residual tumor upon examination within the subcutaneous transplant site (**Figure 6I and S6B-D**). Our observations suggest progressive depletion of the L140P-PUF60 construct over time, implying a growth disadvantage in TNBC cells upon mutation within the RRM1 domain. In conclusion, our investigation into the molecular and functional consequences of a mutation within the PUF60 RRM1 domain strongly substantiate our findings that PUF60 binding at 3’ splice sites is required to sustain TNBC growth and survival in both cells and tumors. Moreover, our findings underscore that TNBC cells lacking PUF60-3’ splice site binding activity show similar gene expression changes to cells with general PUF60 knockdown, confirming that PUF60’s role in regulating cell cycle and DNA metabolism pathways is primarily through its splicing activity at key transcripts required for maintaining the survival and proliferation of TNBC cells.

**Figure 6.**
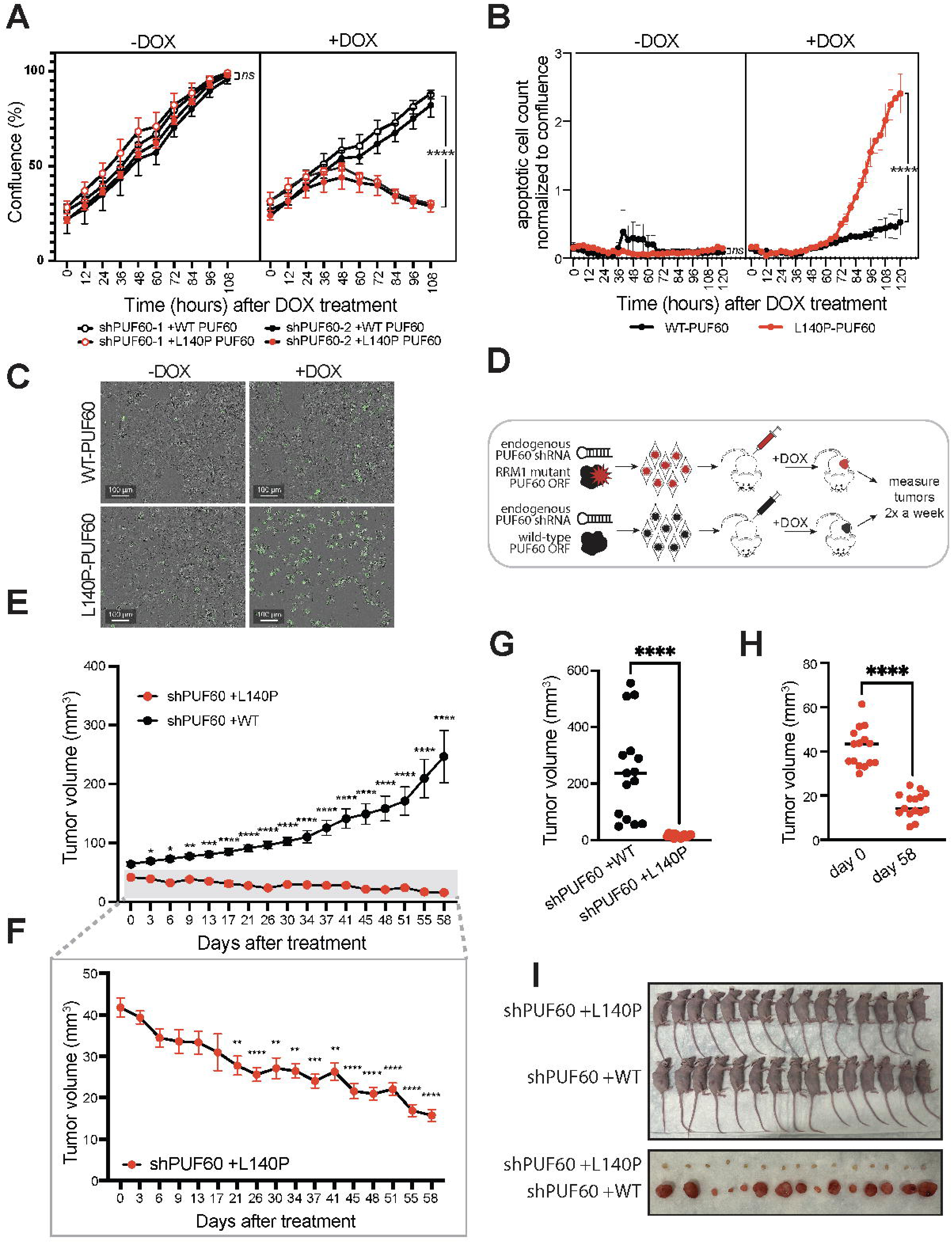
Silencing of PUF60 splice site activity suppresses tumor growth. (A) Cell proliferation after 1μl/ml doxycycline (DOX) treatment (at 0 hours) of WT-PUF60 and L140P-PUF60 MDA-MB-231 cells. Statistical comparisons between WT-PUF60 and L140P-PUF60 quantification at 108 hours. ns: not significant, ****p<0.0001, two-sided T test (B) Caspase 3/7 positive object count per field of view (FOV) normalized to cell confluence per FOV after 1μl/ml doxycycline (DOX) treatment (at 0 hours) of WT-PUF60 and L140P-PUF60 MDA-MB-231 cells. Statistical comparisons between WT-PUF60 and L140P-PUF60 quantification at 120 hours. ns: not significant, ****p<0.0001, two-sided T test (C) Overlayed representative phase and green fluorescent channel images taken at 120 hours for each condition in (B) (D) Schematic describing the generation of DOX-inducible L140P- and WT-PUF60 expressing xenograft growth curves. (E) Average tumor volume over 58 days of DOX-treated mice bearing L140P-PUF60 (red) and WT-PUF60 (black) MDA-MB-231 xenografts. Statistical comparisons between L140P-PUF60 and WT-PUF60 at each time point. *p<0.05, **p<0.01, ***p<0.001, ****p<0.0001, two-way ANOVA with Bonferroni’s test for multiple comparisons. Bars are mean ± SEM; WT-PUF60, n = 15 mice; L140P-PUF60, n = 15 mice. (F) Inset from (E) highlights L140P-PUF60 average tumor volume with reduced y axis range. Statistical comparisons between each time point and initial (day 0) tumor volume, **p<0.01, ***p<0.001, ****p<0.0001, two-way ANOVA with Bonferroni’s test for multiple comparisons. Bars are mean ± SEM; L140P-PUF60, n = 15 mice. (G) Tumor volumes 58 days following DOX induction compared with vehicle controls. ****p<0.0001, paired Student’s t test. (H) Tumor volumes of DOX-induced tumors at initial (day 0) compared with 58 days following treatment. ****p<0.0001, paired Student’s t test. (I) Images of mice and excised tumors 58 days following DOX induction of WT-PU60 and L140P-PUF60 expressing tumors.

## Discussion

Our unbiased *in vivo* functional screen identified 50 candidate RNA-binding protein (RBPs) that contribute to TNBC development within the physiological tumor microenvironment. While these RBPs were implicated in various steps of RNA metabolism, our candidate cohort was significantly enriched for the U2 snRNP spliceosomal complex, which is responsible for intronic branch site identification, and includes RBPs frequently mutated or aberrantly expressed in various cancers, including breast cancer^28, 53, 54^. Consistent with these findings, we identified that the U2-associated splicing factor PUF60 is essential for sustaining TNBC cell and tumor growth. Through integrated eCLIP and knockdown RNA-seq analysis, we revealed that PUF60 drives exon inclusion within transcripts encoding proteins involved in cell cycle and DNA damage pathways. Upon PUF60 depletion, these mRNAs exhibited widespread exon skipping and subsequent destabilization, leading to selective activation of programmed cell death in TNBC. Our observations complement previous work showing that TNBC is especially vulnerable to the inhibition of cell cycle and genomic stability regulators^55–57^. As these pathways collaborate to protect TNBC cells from fatal DNA insults that accompany uncontrolled proliferation rates, our study implicates a role for PUF60 in regulating transcripts that are crucial for TNBC survival^55, 58^. Collectively, our findings reveal how PUF60 modulates the splicing of key cell cycle and DNA repair-encoding transcripts to contribute to cellular survival-associated gene regulation in TNBC.

RBPs are multifunctional proteins and determining their disease-specific regulatory roles remains a significant challenge in cancer biology. Many studies suggest PUF60’s cancer-promoting functions may be RNA-independent^35^ or cytoplasmic^37, 39^. Here, we establish and additional function for PUF60 within the nucleus by disrupting PUF60’s 3’SS direct RNA interactions using a previously characterized RRM1 domain mutation^50^ and identify splicing activity as a key molecular mechanism through which PUF60 contributes to TNBC survival and growth. Our eCLIP analysis of transcripts directly bound by PUF60 reveals a network of candidate targets within proliferation-associated pathways, which safeguard genomic integrity during the mitotic cell cycle. Our multiscale transcriptomics further suggest that PUF60’s oncogenic functions depend on the regulation of several mRNA targets, positioning splicing activity as central to its role in TNBC. As such, our datasets serve as a valuable resource for identifying comprehensive signatures of splicing vulnerabilities in TNBC progression.

Aberrant genomic repair accompanied by chronic replication stress is a dependency shared across many tumors, suggesting that other cancers may require PUF60 for survival. In line with this, several studies have found that PUF60 depletion also induces apoptosis in lung adenocarcinoma^38^ and ovarian cancer^37^, both characterized by replication stress due to high proliferation rates^59^. These findings underscore PUF60’s potential as a broader therapeutic target in cancers with genomic instability.

Our *in vivo* findings underscore the potential of inhibiting PUF60 as a therapeutic strategy to induce TNBC tumor regression. While several transcriptional and translational modulators have been developed, few spliceosomal inhibitors have been thoroughly characterized, and none are specific to PUF60^60–63^. However, significant progress has been made toward developing compounds exhibiting anti-cancer properties that inhibit the U2-associated SF3b1 protein complex^62–65^. Cancers driven by aberrant spliceosome activity are highly sensitive to SF3b inhibition^66–68^, underscoring the potential of selective and potent small-molecule splicing modulators for the treatment of aggressive cancers that lack effective targeted therapies, such as TNBC. Although PUF60-specific inhibitors have yet to be characterized, small molecules that block interactions between U2AF homology motifs (UHM) and UHM ligand motifs (ULM)—domains found in various proteins including U2-associated factors such as PUF60—have been developed^69, 70^. Targeting protein-protein interactions between PUF60 and spliceosomal factors could be advantageous in TNBC, but the presence of UHM and ULM domains in non-spliceosomal proteins, such as kinases, raises concerns about potential off-target effects from these inhibitors^71^. To mitigate these challenges, we propose exploring PUF60-specific inhibitors as a therapeutic strategy for TNBC; however, future studies are essential to evaluate the safety of PUF60 inhibition. Alternatively, splice-switching antisense oligonucleotides (ASOs) that bind to pre-mRNA and block splicing machinery interactions could reverse specific downstream splicing events and reduce potential toxicity from general PUF60 inhibition^72^. The specific downstream AS events that only contribute to apoptosis in PUF60-depleted TNBC cells have not yet to be determined. Since an individual splicing variant is unlikely to account for the entirety of PUF60’s role, future investigations should examine the combinatorial impact of multiple splicing variants on TNBC survival. Our comprehensive study provides a molecular framework for identifying critical splicing events that could be leveraged to disrupt genomic maintenance and cell cycle processes that safeguard TNBC against replication stress.

## Key Resources Table

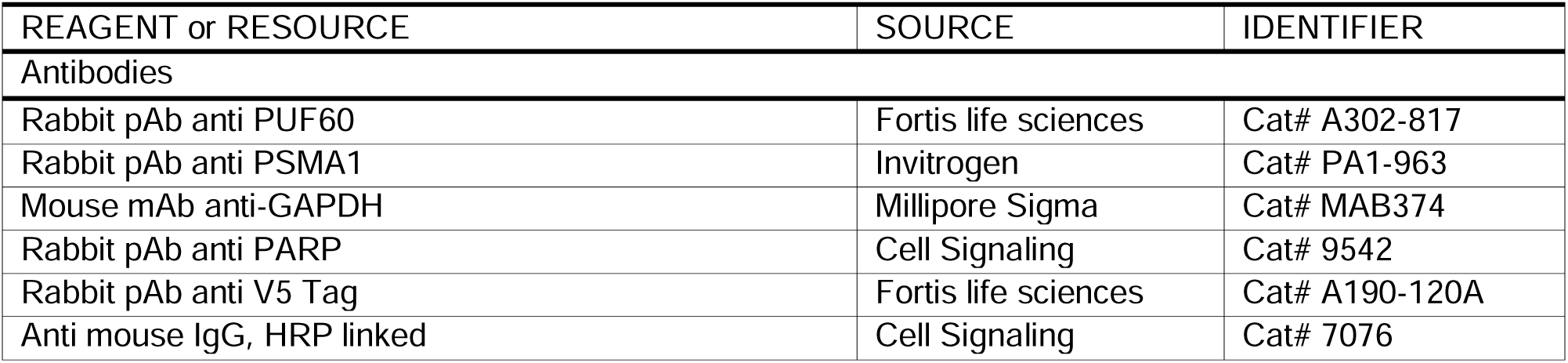
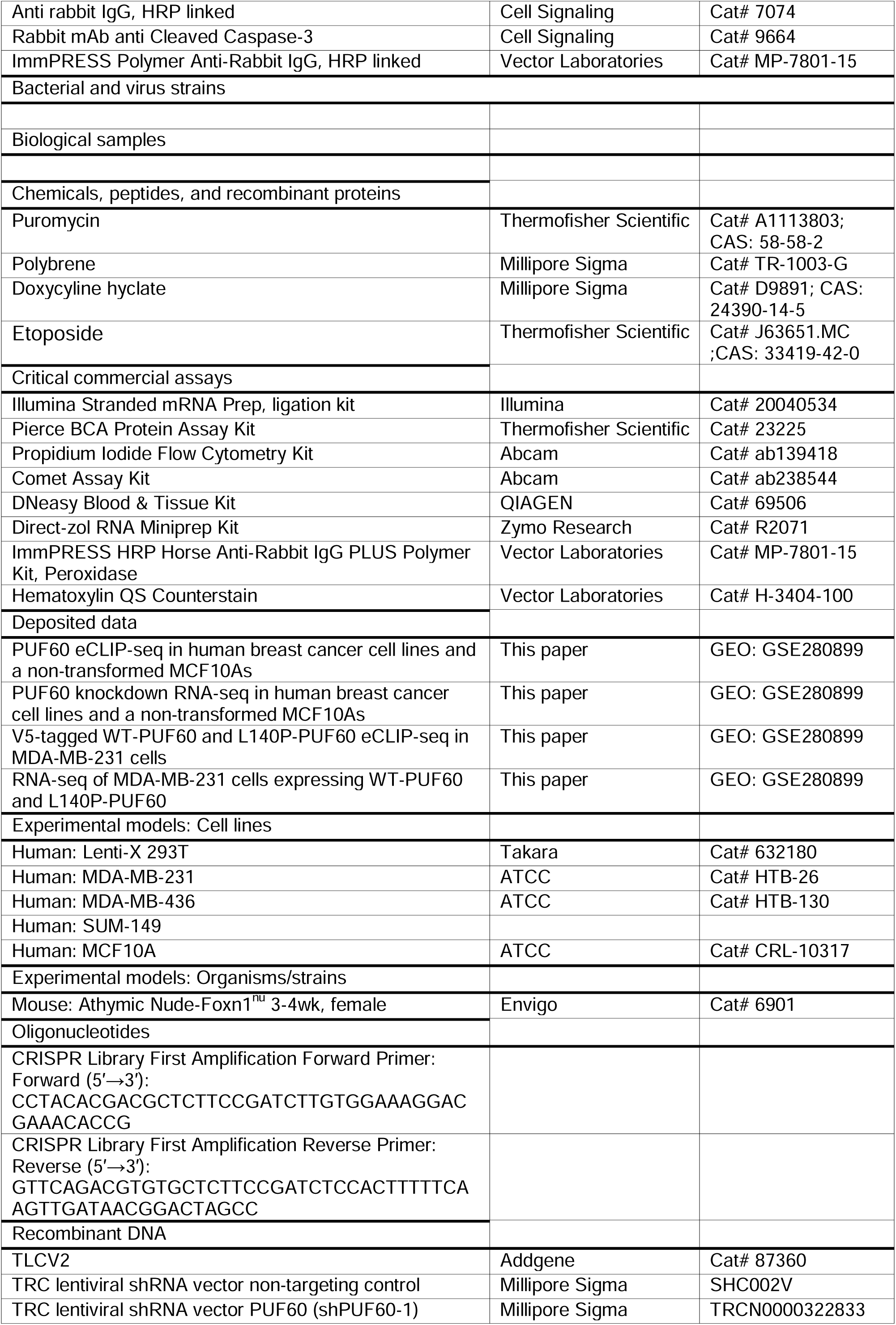
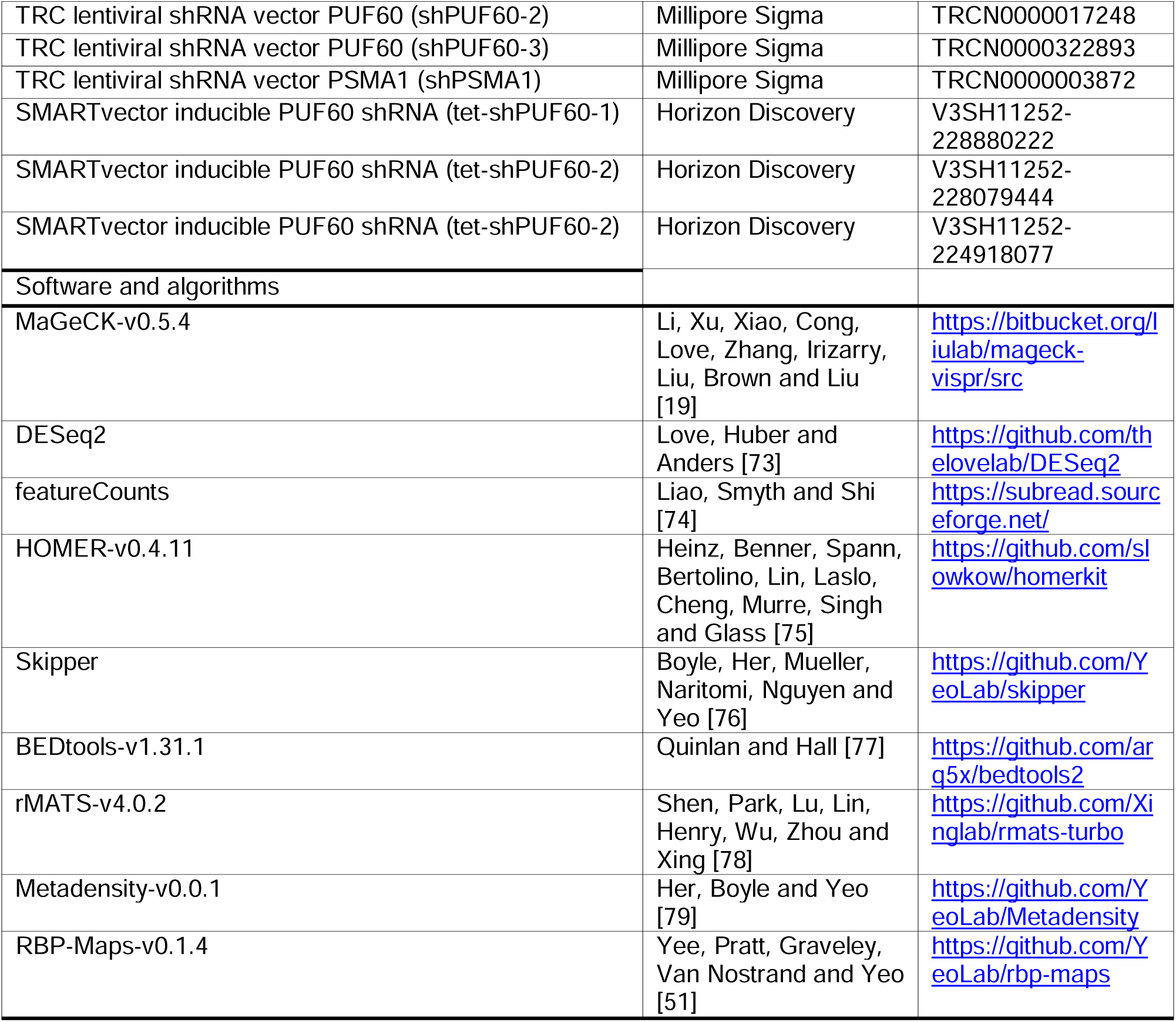

## STAR Methods

### Experimental model and subject details

#### Cell culture

MCF10A (female) immortalized cells were maintained in Human Mammary Epithelial Cell Basal Medium supplemented with 0.4% bovine pituitary extract, 0.01 μg/ml recombinant human insulin-like growth factor, 0.5 μg/ml hydrocortisone, and 3 ng/ml human epidermal growth factor. MDA-MB-231 (female) immortalized cells were maintained in DMEM supplemented with 10% FBS. MDA-MB-436 (female) immortalized cells were maintained in DMEM supplemented with 10% FBS and 10 μg/ml recombinant human insulin. SUM-149 (female) immortalized cell lines were maintained in Ham’s F-13 Nutrient Mix supplemented with 5% FBS, 1 μg/ml hydrocortisone, and 5 μg/ml recombinant human insulin. All cells were passaged every 2 or 3 days with TrypLE Express Enzyme and maintained in a humidified incubator at 37°C with 5% CO_2_. Cells were routinely tested for mycoplasma contamination with a MycoAlert mycoplasma test kit (Lonza) and were found negative for mycoplasma.

#### Animal Studies

Animal protocols were approved by the University of California, San Diego IACUC under protocol S12099. Athymic Nude-Foxn1^nu^ were purchased at 6-weeks of age from Envigo International Holdings, Inc.

### Method details

#### CRISPR plasmid library preparation

A subset of sgRNA sequences from our previously developed CRISPR-Cas9 lentiviral library ^80^ were ordered as a pool of equal molar oligonucleotides (Supplementary Table). The lentiCRISPR RBP plasmid library was cloned using previously reported methods ^81^. To construct a doxycycline-inducible library, the TLCV2 backbone ^18^ was digested with BsmBI and PCR amplified sgRNA oligonucleotide inserts were Gibson assembled. To maintain 300x library complexity, plasmid products were amplified using parallel electroporations. Electroporated bacteria were spread on carbenicillin agar plates (24.5cm x 24.5cm) and grown for 16-18 hours at 32°C. One day later, colonies were scraped, collected, and maxiprepped.

#### Lentivirus production and purification

HEK293xT cells were plated on 15 cm plates at 70% confluency the day before transfection. 1 hour before transfection, media was removed and replaced with 15ml of pre-warmed OptiMEM. Transfections were carried out using 62.5 μl Lipofectamine 3000, 125 μl of P3000 Enhancer, 12.5 μg lentiCRISPR plasmid library, 6.25 μg pMD.2g, and 9.375 μg psPAX2. 8 hours post-transfection, media was changed to DMEM supplemented with 10% FBS. After 48 hours, the media supernatant was filtered through a 0.45 μm nitrocellulose membrane. The virus was ultracentrifuged at 24,000 rpm for 2 hours at 4°C and resuspended overnight in PBS at 4°C.

#### Multiplicity of infection

To determine the volume of virus required for an MOI of 0.3 for each cell type, virus was titrated across a 6-well plate with 1 million cells per well in medium containing 8 μg/mL polybrene. After 48 hours, the medium was replaced with DMEM supplemented with 10% FBS and 2 μg/mL puromycin. Three days later, the number of surviving cells was counted to establish the MOI based on the virus volume that resulted in 30% cell survival.

#### In vitro RBP CRISPR screen

MCF10A, MDA-MB-231, MDA-MB-436, SUM-149 cells were infected with RBP CRISPR library lentivirus at 0.3 MOI in medium containing 8 μg/mL polybrene. After 24 hours, media without polybrene was replaced. Following an additional 48 hours, media containing 2 μg/mL puromycin was replaced and maintained on for 4 days to select for infected cells. Cells expressing the RBP CRISPR library were then expanded plating 4 million cells per 15cm plate, with a total of 2 plates per replicate. Once plate from each replicate was treated with 1 μg/mL DOX. The cells were cultured for 14 days, with media and DOX replaced every 2 days and cell splitting every 4 days. For each condition and replicate, 4 million cells were harvested and snap frozen.

#### *In vivo* RBP CRISPR screen

MDA-MB-231s, MDA-MB-436s, and SUM-149s expressing the RBP CRISPR library were expanded and subcutaneously transplanted (3 x 10^6^ cells per mouse to maintain 200x library complexity) into the right flank of athymic nude mice (female, 6 weeks old). Following sufficient tumor engraftment (tumor volume of ∼300 mm^3^), mice were randomized and maintained 5% sucrose water (5 mice/cell line, -DOX) or 5% sucrose water with 2mg/mL DOX (Sigma Aldrich; D9891) (5 mice/cell line, +DOX) for 14 days. Tumors were washed with sterile PBS and minced with a sterilized blade into 2-3mm cubes. Tumor pieces were dissociated in enzyme medium (∼0.1cm^3^ tumor/∼2mL), shaking for 30 minutes at 37°C. The enzyme medium consisted of 10x enzyme mix (Collagenase IV, 1g/100mL HBSS; (Sigma; #C-5138), Hyaluronidase, 100mg/100mL HBSS; (Sigma, #H-6254), and Deoxyribonuclease, 20,000 U/100 mL HBSS; (Sigma; #D-5025)) in RPMI 1640 medium with 10% Pen/Strep. 4×10^6^ cells were spun down for each replicate and snap frozen

#### Bulk sgRNA library preparation for *in vitro* and *in vivo* RBP CRISPR screens

DNA libraries were prepared using a targeted-enrichment approach as previously described ^80^. Briefly, genomic DNA (gDNA) was extracted from pellets of 4 million cells using DNeasy Blood & Tissue kit (Qiagen; 69504), then sonicated to ∼1000bp by Biorupter. The average fragment size was assessed with genomic DNA ScreenTapes on the Agilent Tapestation (Agilent, 5067-5365). Fragments containing sgRNAs were isolated with biotinylated RNA probes that target a conserved region on the TLCV2 backbone, and these were captured with streptavidin beads. Isolated gDNA fragments were purified using the DNA Clean and Concentrator-5 kit (Zymo Research; 11-303C). PCR was then performed with primers flanking the sgRNA:

Forward (5′→3′): CCTACACGACGCTCTTCCGATCTTGTGGAAAGGACGAAACACCG;

Reverse (5′→3′): GTTCAGACGTGTGCTCTTCCGATCTCCACTTTTTCAAGTTGATAACGGACTAGCC

Followed by a second amplification with Illumina sequencing primers. Library quality was evaluated using an Agilent D1000 Screen Tape on the Agilent Tapestation (Agilent; 5067-5582) and then sequenced to 6M reads per library on the Hi-Seq4000 in paired-end 55bp mode.

#### Statistical analysis of in vivo and in vitro RBP CRISPR screens

Reads were aligned to the RBP library and the MaGeCK version 0.5.4 software package was used to statistically identify RBP candidates ^19^. RBP candidates with a Wald P value < 0.05 and beta score < 0 were considered candidate essential RBPs.

#### TCGA and GTEx data description

Publicly available datasets from The Cancer Genome Atlas Breast Invasive Carcinoma (TCGA-BRCA) and The Genotype-Tissue Expression (GTEx) was directly downloaded from the UCSC Xena portal at https://xena.ucsc.edu/ ^82^. For gene expression data, we used mRNA expression z-scores calculated by RNA-seq by Expectation Maximum (RSEM) and categorized the data by samples from normal breast tissue (normal) and basal clinical tumor subtype (basal). For survival data, the Kaplan-Meier probability was determined using the UCSC Xena portal using clinical overall survival status amongst TCGA-BRCA patients by grouping the 1^st^ and 4^th^ quartiles of PUF60 copy number variations (CNVs) determined by the Log_2_(Tumor/normal).

#### Knock-down experiments

Cells were transduced with the TRC lentiviral shRNA vector non-targeting control (NTC; Millipore Sigma; SHC002), and TRC lentiviral shRNA vector PUF60 (shPUF60-1; Millipore Sigma; TRCN0000322833), (shPUF60-2; Millipore Sigma; TRCN0000017248), (shPUF60-3; Millipore Sigma; TRCN0000322893), or TRC lentiviral shRNA vector PSMA1 (shPSMA1; Millipore Sigma; TRCN0000003872) for 48 hours before treatment with 2 μg/mL puromycin. Cells were analyzed 4 days after the addition of lentivirus for all assays unless noted otherwise.

#### Western blot

Cells were lysed with RIPA buffer (Sigma Aldrich; R0278) containing protease inhibitor (Thermofisher, A32953). Protein lysates were centrifuged to pellet and remove insoluble material and were then quantified using the Pierce BCA Kit. Protein lysates were run on a 4-12% NuPAGE Bis-Tris gel and transferred to a polyvinylidene fluoride (PVDF) membrane. Membranes were blocked in Tris-Buffered Saline containing Tween 20 (TBST) with 5% milk for 20 minutes and probed overnight at 4C with the following primary antibodies: Rabbit pAb anti-PUF60 (Fortis life sciences, A302-817), Rabbit pAb anti-PSMA1 (Invitrogen, PA1-963), Mouse mAb anti-GAPDH (Millipore Sigma, MAB374), Rabbit pAb anti PARP (Cell Signaling, 9542), and (Rabbit pAb anti V5 Tag, Fortis life sciences, A190-120A). Membranes were washed 3 times for 5 minutes with TBST and then probed for 1 hour at room temp in TBST containing 5% milk with the following secondary antibodies diluted 1:5000: Anti mouse IgG, HRP linked (Cell Signaling, 7076), and Anti rabbit IgG, HRP linked Cell Signaling, 7074).Membranes were washed 3 times for 5 minutes with TBST and developed using Thermo Pierce ECL detection kits on an Azure Western Blot Imaging System.

#### Plasmid construction and generation of L140P-PUF60 and WT-PUF60 stable cell lines

To generate L140P-PUF60 and WT-PUF60 stable cell lines, the inducible Lentiviral CRISPRi expression backbone (addgene #167935) was digested at XHO1 and BAMH1 sites to remove the KRAB, dCas9, and DHFR domains positioned immediately downstream of the Tet operator sequence. gBlocks (IDT) containing codon-optimized PUF60 ORFs with (L140P-PUF60) and without (WT-PUF60) the L140P mutation, a C-terminal V5 tag and 25nt complimentary sequences were assembled with the digested Lentiviral backbone (immediately downstream of the Tet operator sequence) in a two-fragment Gibson Assembly reaction. To generate lentiviral particles for Inducible L140P-PUF60, WT-PUF60 and SMARTvector inducible PUF60 shRNAs (tet-shPUF60-1; Horizon Discovery; V3SH11252-228880222), (tet-shPUF60-2; Horizon Discovery; V3SH11252-228079444), (tet-shPUF60-3; Horizon Discovery; V3SH11252-224918077), we seeded HEK293T cells on 10cm plates at 70% confluency, 24 hours prior to transfection. Transfections were performed using 45 μl Lipofectamine 3000, 60 μl of P3000 Enhancer, 7 μg transgene, 3.6 μg pMD.2g, and 6.6 μg psPAX2. 8 hours following transfection, media was changed to DMEM + 10% FBS. 48 hours following transfection, the media supernatant was filtered through a 0.45 μm low protein binding membrane. The virus was concentrated with Lenti-X Concentrator (Takara) and centrifuged and resuspended according to manufacturer instructions. To make tet-shPUF60 cell lines, MDA-MB-231s were individually transduced with the inducible PUF60 shRNAs for 48 hours before treatment with 2 μg/mL puromycin. To make L140P-PUF60 and WT-PUF60 cell lines, stable tet-shPUF60-1 and tet-shPUF60-2 MDA-MB-231 cell lines were then transduced with L140P-PUF60 and WT-PUF60 lentivirus and sorted for the top 10% of RFP-expressing cells on a BD Influx Cell Sorter.

#### eCLIP-seq library preparation

Experiments were performed as previously outlined ^41^ with biological duplicates. In brief, 20M cells were UV-crosslinked at 400 mJ/cm^2^ constant energy, lysed, and sonicated by BioRuptor. Lysates were treated with RNase 1 to fragment RNA and then protein-RNA complexes were immunoprecipitated with PUF60 antibody (Rabbit pAb anti-PUF60, Fortis life sciences, A302-818) or V5 antibody (Rabbit pAb anti V5 Tag, Fortis life sciences, A190-120A) pre-coupled to Sheep anti-rabbit Dynabeads. Paired inputs (2% of lysate) were first removed from each sample. IP samples were stringently washed, and RNA from all samples was dephosphorylated with FastAP (Thermofisher) and T4 RNA ligase (NEB), followed by on-bead ligation of barcoded RNA adapters to the 3’ end of RNA in sample (NEB). RNA-protein complexes were run on 4-12% NuPAGE Bis-Tris gels and transferred to nitrocellulose membranes where the RNA in the region 60kDa – 140kDa was excised from the membrane and proteinase K treated (NEB) to release RNA. Input samples were dephosphorylated with FastAP (Thermofisher) and T4 PNK and a 3’RNA adapter was ligated with T4 RNA ligase (NEB) so synchronize input and IP samples. RNA was reverse transcribed using Superscript III (Thermofisher), followed by ExoSAP-IT (Applied Biosystems) to remove excess primers. qPCR was used to determine the appropriate number of PCR cycles for library amplification, followed by amplification with Q5 (NEB; M0492S) and QC using an Agilent D1000 Screen Tape (Agilent). Libraries were sequenced to 20M reads on the HiSeq 2000, 2500, or 4000 platform.

#### Computational analysis of eCLIP data

Reproducible eCLIP peaks were called using the latest release of the Skipper pipeline, available on GitHub (https://github.com/YeoLab/skipper) ^76^. In brief, Skipper trims and aligns reads, determines counts per window, and compares enrichment in IP samples against input samples. Enriched elements are then considered reproducible if they meet a 20% false discovery threshold in both replicates for each sample. Only those with a significant FDR less than 0.05 and enrichment odds ratio greater than 3 were included in further analysis. For V5-tagged eCLIPS, nonspecific V5 antibody binding in wild-type MDA-MB-231s was removed from the analysis of L140P-PUF60 and WT-PUF60 eCLIPS using BEDTools version 1.31.1^77^. Motifs were analyzed using HOMER version 4.11^75^, and metagene plots were generated using Metadensity version 0.0.1^79^. Overlapping windows amongst cell types were determined using BEDTools version 1.31.1^77^, and splicing maps were generating using RBP-Maps version 0.1.4^51^.

#### RNA-seq library preparation

To assess the impact of PUF60 knockdown on both normal and TNBC cell lines, total mRNA was isolated from 3 biological replicates of non-targeting controls and 3 biological replicates for each PUF60-targeting shRNA (shPUF60-1, shPUF60-2, and shPUF60-3) in MCF10A, MDA-MB-231, MDA-MB-436, and SUM-149 cell lines. To evaluate the effect of the L140P mutation on PUF60 splicing activity, total mRNA was extracted from 3 wild-type PUF60 and 3 L140P mutant biological replicates in MDA-MB-231 cells. RNA quality was assessed using an Agilent RNA ScreenTape (Agilent). A total of 500 ng RNA was rRNA-depleted, and library preparation was carried out with the Stranded mRNA Prep Ligation Kit (Illumina). Libraries were quality controlled using an Agilent D1000 Screen Tape (Agilent). Sequencing was performed on HiSeq 2000, 2500, or 4000 platforms, collecting 60 million 150bp paired-end reads per sample.

#### Computational analysis of integrated RNA-seq and eCLIP data

Adapters trimming and read mapping to the human genome build hg38 was performed using STAR version 2.4.0. Differential expression analysis for genes with a minimum TPM of 10 in any sample was conducted with DEseq2 version 1.22.1 ^73^. Differential AS events were analyzed using rMATS version 4.0.2 ^78^, with only differential splicing events with a sum of 30 reads across all conditions considered for downstream analysis. Significant differentially splicing was defined by an absolute inclusion-level difference greater than 5% and an false discovery rate (FDR) of less than 5%. Exon splicing events containing an adjacent upstream 3’SS eCLIP window were determined using BEDtools version 1.31.1 ^77^.

#### Gene Ontology (GO) analysis

Gene ontology enrichment was analyzed using Metascape version 3.5 ^83^, with background genes consisting of those expressed with a TPM greater than 10 in each cell line for the integrated eCLIP and RNAseq analysis. For the CRISPR screen candidate ontology analysis, the entire cohort of RBPs from the Lenti CRISPR library utilized as the background list.

#### Cell cycle analysis

MDA-MB-231 and MCF10A cells were transduced with NTC, shPUF60-1, shPUF60-2, or shPUF60-3 virus and selected for 2 days with 2 μg/mL puromycin (Thermofisher Scientific, A1113803). 5 days after transduction, cells were stained with zombie violet dye from the Zombie Violet Fixable Viability Kit (BioLegend, 423113) according to manufacturer’s instructions. Then, cells were fixed in 66% ice cold ethanol and stained with propidium iodide using the Propidium Iodide Flow Cytometry Kit (Abcam, ab139418) according to manufacturers instructions. Cells were analyzed by flow cytometry using the BDSLRFortessa under the Pacific blue (Zombie Violet) and PE-Texas Red (Propidium Iodide) channels. Analysis and gating were performed using FlowJo.

#### Comet Assay

The comet assay was performed according to the manufacturer’s instructions (Abcam, ab238544). Briefly, MDA-MB-231 and MCF10A cells were seeded in 12-well plates 4 days after transduction with either non-targeting control NTC, shPUF60-1, shPUF60-2, or shPUF60-3 virus. Cells expressing shNTC were either treated with either 40 μM Etoposide or DMSO vehicle control for 4 hours before collection by gentle scraping. A total of 1,000 cells in 10 μl were mixed with 60ul agarose and transferred onto comet slides. The agarose-cell suspension was allowed to solidify followed by lysis at 4C for 1 hour. Electrophoresis was performed with cold TBE buffer for 15 minutes at 60 volts. The comet slides were rinsed in H_2_O 3 times and then fixed with 70% cold ethanol for 5 min. Air-dried slides were then stained with Vista Green DNA Dye and viewed on a fluorescence Keyence microscope. Percentage of DNA in the tail, tail length, and tail moment were assessed using the ImageJ plugin OpenComet, with 50-100 nuclei measured per condition.

#### PUF60 knockdown *in vivo* tumorgenicity assays

To determine the effect of PUF60 knockdown on tumor growth, tet-shRNA-1 and tet-shRNA-2 MDA-MB-231 cells selected with 2 μg/mL puromycin were subcutaneously transplanted (3×10^6^ cells per mouse) into 20 athymic nude mice (female, 6 weeks old) per shRNA. Mice were randomized and half were maintained on 5% sucrose water (-DOX) or 5% sucrose water with 2mg/mL dox (Sigma Aldrich; D9891) (+DOX) 14 days post-transplantation. Twice each week, tumors were measured with calipers and mouse body weight was recorded. RFP-positive, Dox-treated mice were imaged in an IVIS spectrum *in vivo* imaging system (Revvity) on day 3 (initial) and day 46 (final) of dox treatment. Tumors were harvested after reaching 1000mm^3^ on average, followed by snap freezing for protein extraction or fixation in 4% paraformaldehyde for IHC analysis.

#### Immunohistochemistry

Tumor samples were fixed in 4% paraformaldehyde and then paraffin-embedded. Sections of 5 μM thickness were prepared using a microtome by the Moores Cancer Center Histology Core. Sections were deparaffinized in Histoclear and rehydrated through a graded alcohol series. Antigen retrieval was performed with citrate buffer. Tissues were stained with primary antibody against rabbit mAB anti Cleaved Caspase-3 (Cell Signaling #9664; 1:150) and subsequently with a secondary horse anti-rabbit IgG using ImmPRESS kit (Vector Laboratories #MP-7801-15). Slides were stained with diaminobenzidine (DAB) chromogen (Vector Laboratories #SK-4105) followed by Hematoxylin QS (Vector Laboratories #H-3404-100) counterstaining. Images were acquired on the Revolve microscope (Echo Laboratories) using 10X objectives for quantification and 17X for figure representation. Quantification of Cleaved Caspase-3-positive cells was performed with QuPath-0.5.1 (Analyze - Cell detection - Positive cell detection), analyzing 20-35 images per tissue section from each of 9 DOX-treated tumors and 10 vehicle control tumors. The mean percentage of cleaved caspase 3-positive cells per tumor was then used to assess significant differences in cleaved caspase 3 signal between DOX-treated and vehicle control tumors.

#### L140P-PUF60 and WT-PUF60 in vivo tumorgenicity assays

To determine the effect of L140P PUF60 mutation on tumor growth, RFP-sorted L140P-PUF60 and WT-PUF60 MDA-MB-231 cells were subcutaneously transplanted (3×10^6^ cells per mouse) into athymic nude mice (female, 6 weeks old) (15 mice for L140P-PUF60 and 15 mice WT-PUF60). All mice were maintained on 5% sucrose water with 2mg/mL dox (Sigma Aldrich; D9891) (+DOX) 14 days post-transplantation. Twice each week, tumors were measured with calipers and mouse body weight was recorded. RFP-positive, Dox-treated mice were imaged in an IVIS spectrum *in vivo* imaging system (Revvity) on day 0 (initial) and day 58 (final) of dox treatment. Tumors were harvested after reaching 1000mm^3^ on average, followed by snap freezing for protein extraction.

#### Time lapse microscopy

To measure confluence over time, L140P-PUF60 and WT-PUF60 MDA-MB-231s were seeded at 10k cells in Incucyte ImageLock plates (Essen BioSciences; 4379) 24 hours prior to imaging. Plates were loaded into the Incucyte and imaged at 10x magnification for 108 hours every 12 hours. Phase images were analyzed using the Incucyte ZOOM Basic Analyzer.

To quantify apoptotic cells, L140P-PUF60 and WT-PUF60 MDA-MB-231 cells were seeded at a density of 900 cells per well in 384 well plates 24 hours before imaging. Fresh media supplemented with Caspase 3/7 live-cell dye (Sartorius, 4440) was added just prior to imaging. The plates were then loaded into the CellcyteX imaging system from Cytena and imaged at 10x magnification over a period of 120 hours, with images captured every 4 hours. Phase contrast and green fluorescent channel images were analyzed using the Cellcyte Studio to assess cell confluence and measure fluorescent object count per field of view (FOV). Caspase 3/7 FOV values were then normalized to total cell confluence per FOV.

#### Quantification and statistical analysis

Researchers overseeing and measuring individual tumor xenografts were not blinded to the experimental conditions. Mice were assigned to experimental groups through simple randomization. All animal studies were conducted in accordance with institutional and national animal welfare regulations. Power analysis was employed to determine the sample size necessary for detecting significant changes in tumor size. Details of statistical tests, including the test used, exact sample sizes, and precision measures, are provided in the figure legends. A p-value of less than 0.05 was considered significant, with specific p-values reported in the figures or their legends.

#### Resource Availability

Further information and request for resources and reagents should be directed to the lead contact, Gene W. Yeo. RBP CRISPR plasmid library ***will be*** available on Addgene (***XXXX***). CRISPR screening, eCLIP-sequencing, and RNA-seq datasets generated for this study are deposited under GEO accession number GSE280899.

## Acknowledgements

We would like to thank Yeo laboratory members K. Rhine, K. Rothamel, and W. Brothers for critical reading of the manuscript and B. A. Yee and H. L. Her for providing guidance for the computational analysis of eCLIP and RNAseq datasets in this manuscript. We are grateful to Dr. Igor Vorechovsky for kindly providing constructs for generating wild-type and L140P PUF60 cell lines. We thank the Sanford Consortium Human Embryonic Stem Cell Core for providing us use of their instruments and cell-sorting services (1S10OD025060). This publication includes data generated using an Illumina NovaSeq 6000 at UC San Diego provided by the IHM Genomics Center (S10OD026929). Results published here are based, in part, upon data generated by the TCGA Research Network and the Genotype-Tissue Expression project. A.T.T. is supported by the Cancer Systems Biology Training Program (U54 CA209891) and the Biology, Informatics, and Omics Training Program (T32CA067754). O.M. is supported by a Gruss-Lipper postdoctoral fellowship. M.P. is supported by the Ruth L. Kirschstein F32 National Research Service Award (F32 HL143978) from the NIH. The authors acknowledge support from the National Institutes of Health (U24 HG009889, RF1 MH126719, R01 HG011864, R01 HG004659 to G.W.Y., and a subaward to G.W.Y. from U24 HG011735). This project has been made possible in part by grant 2023-332369 from the Chan Zuckerberg Initiative DAF, an advised fund of the Silicon Valley Community Foundation. This work was supported by the V Foundation for Cancer Research (V2024-008 to C.E.A) and by the American

Cancer Society IRG Grant # IRG-19-230-48-IRG and UC San Diego Moores Cancer Center, Specialized Cancer Center Support Grant NIH/NCI P30CA023100 to C.E.A.

## Author Contributions

Conceptualization, A.T.T., J.M.E., and G.W.Y.; methodology, A.T.T., J.M.E., O.M., and D.K.; formal analysis, A.T.T and J.M.E; investigation, A.T.T., J.M.E., C.J.Z., V.P., J.T.N., G.G.N., M.P., J.S., Y.Z., and F.E.T.; data curation, A.T.T., J.M.E., C.J.Z. and V.P.; writing – original draft, A.T.T.; writing – review & editing, A.T.T., O.M., M.P., C.E.A. and G.W.Y.; visualization, A.T.T. and J.M.E.; supervision, D.S.K. and G.W.Y.; funding acquisition, D.S.K., C.E.A. and G.W.Y.

---

**Figure S1.**
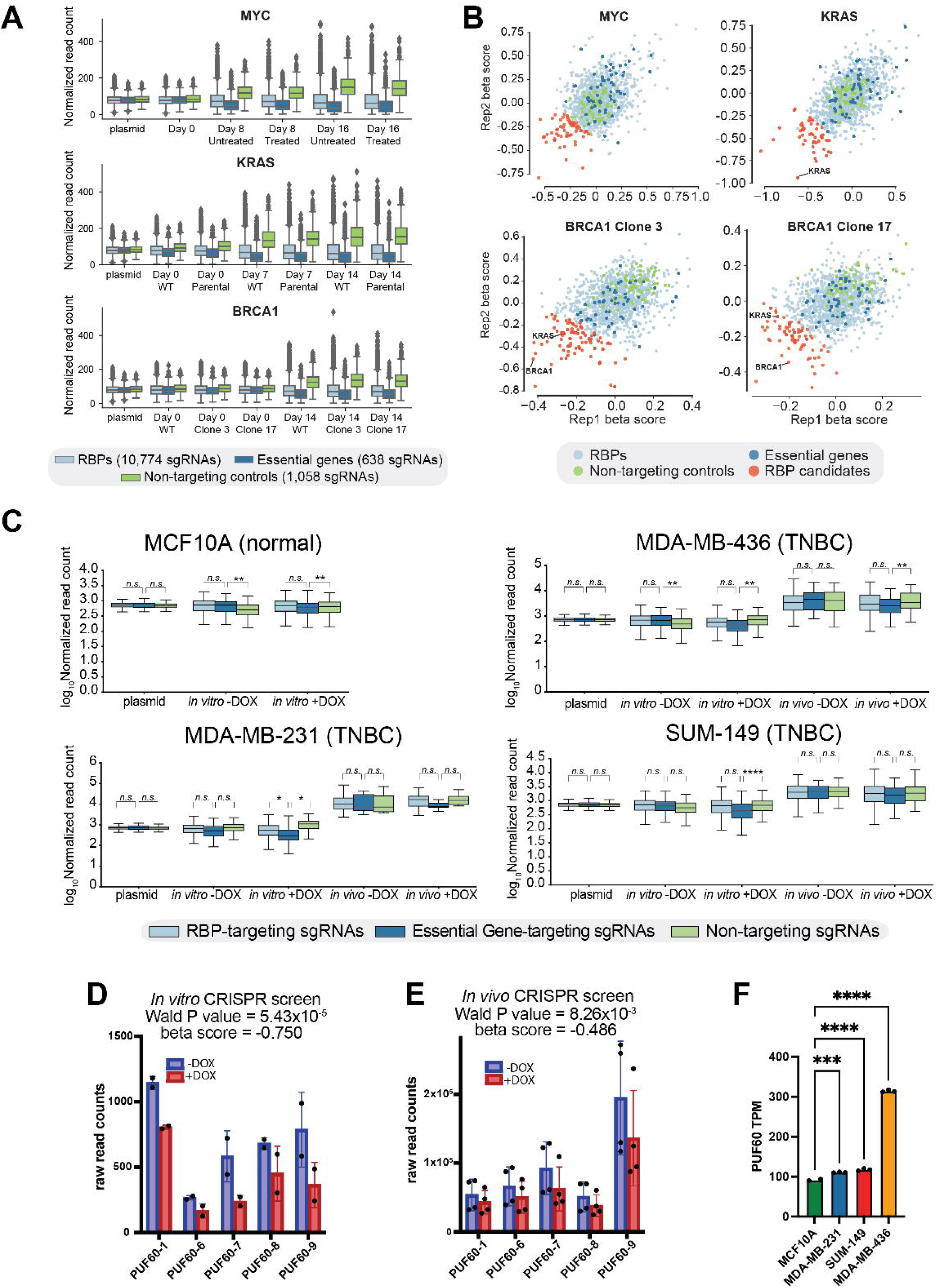
CRISPR screening in isogenic cell lines and TNBC-derived cell lines identifies PUF60 as an essential RBP in TNBC, related to figure 1. (A) Boxplots comparing normalized sgRNA abundance in MYC amplification, KRAS G13D gain-of-function (GoF), and BRCA1 loss-of-function (LoF) isogenic cell lines. Each condition is representative of 2 independent replicates. (B) Comparison of β score replicates on day 16 (MYC) or day 14 (BRCA1 and KRAS) for RBPs (light blue), non-targeting controls (green), essential genes (dark blue) and RBP candidates (red). (C) Boxplots comparing normalized sgRNA abundance following doxycycline (DOX)-induced Cas9 expression *in vitro* and *in vivo*. *In vivo* conditions are representative of 5 independent replicates and in vitro conditions are representative of 2 independent replicates. *p<0.05, **p<0.01, ****p<0.0001, one-way ANOVA with Dunnett’s test for multiple comparisons. (D) Bar plot illustrating raw abundance of sgRNAs targeting PUF60 following DOX-induced Cas9 expression *in vitro*. MAGeCK Wald test. (E) Bar plot illustrating raw abundance of sgRNAs targeting PUF60 following DOX-induced Cas9 expression *in vivo*. MAGeCK Wald test. (F) Bar plot illustrating PUF60 transcripts per million (TPM) among TNBC and normal mammary epithelial cell lines. ***p<0.001, ****p<0.0001, one-way ANOVA with Dunnett’s test for multiple comparisons.

**Figure S2.**
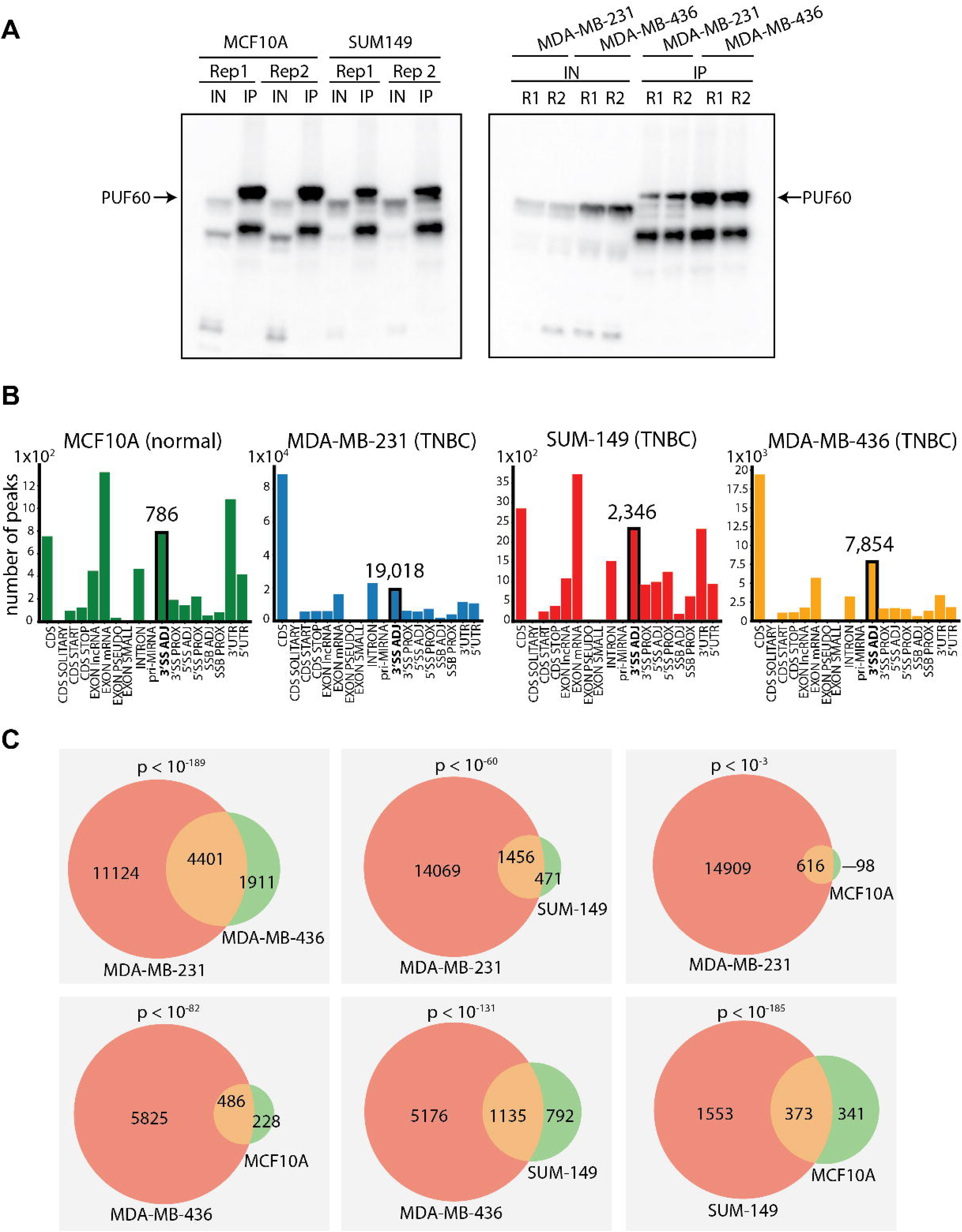
Quality control of PUF60 eCLIP and cell line comparisons of 3’ splice site targets, related to figure 2. (A) Western blots of eCLIP immunoprecipitations for PUF60 in MCF10A, MDA-MB-231, MDA-MB-436, and SUM-149 cells. (B) Bar plots summarizing gene region-specific distribution of reproducible eCLIP peaks. Peaks adjacent to 3’ splice sites are outlined in black. (C) Venn diagram comparisons of reproducible 3’SS PUF60 eCLIP peaks among all 4 cell lines. Fisher’s exact test.

**Figure S3.**
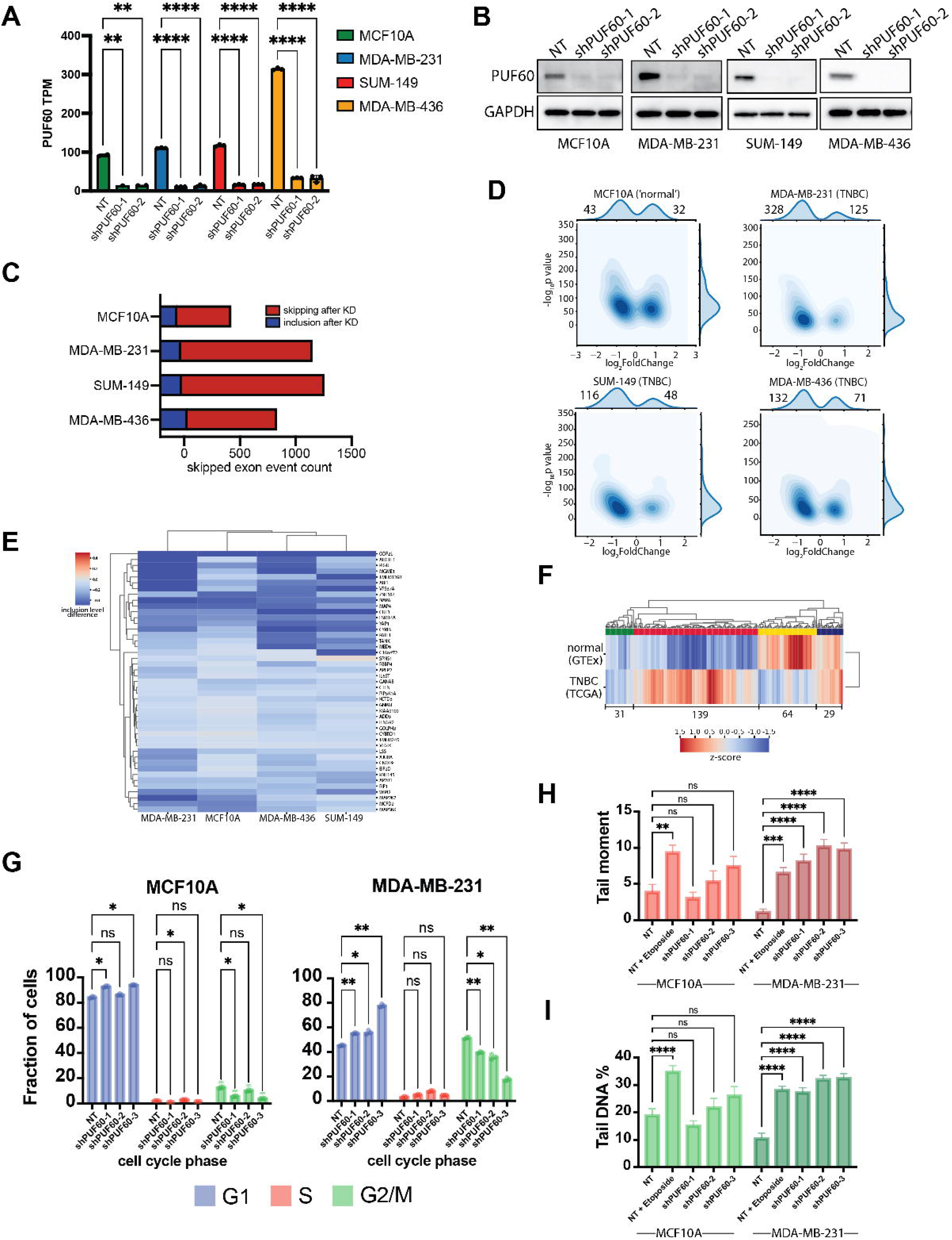
KD RNA-sequencing reveals exon skipping events modulated by PUF60 in TNBC and normal cells, related to figure 3. (A) PUF60 TPM following shRNA knockdown as measured from aligned RNA-seq data, **p<0.01, ****p<0.0001, one-way ANOVA with Dunnett’s test for multiple comparisons. (B) Western blot analysis of cell lysates from non-targeting control (NTC), shPSMA1, and shPUF60 TNBC and normal mammary epithelial cell lines. (C) Stacked bar plot of differentially spliced exons after shRNA-mediated knockdown of PUF60 as identified by rMATS analysis. Differential exon inclusion events (blue) have an inclusion level difference > 0.05 and differential exon exclusion events (red) have an inclusion level of < −0.05. All differentially spliced exons have a false discovery rate (FDR) < 0.01 as calculated by the rMATS log likelihood ratio. (D) Jointplots illustrating the expression of genes containing KD-sensitive exons bound by PUF60 as identified by DESeq2 analysis. (E) Hierarchical cluster map of inclusion levels differences from exons within the red cluster in (Fig. 3D). (F) Hierarchical cluster map summarizing mRNA expression levels Z scores of genes within grey and blue clusters from (**F**) between basal-like breast cancer (TCGA data portal) ^24^ and normal breast tissues (GTEx comprehensive tissue resource) ^25^. (G) Quantification of propidium iodide (PI) staining in MCF10As and MDA-MB-231s expressing non-targeting control shRNA and PUF60-targeting shRNAs. *p<0.05, **p<0.01, two-way ANOVA with Holm-Sidak’s test for multiple comparisons. (H) Bar plot illustrating quantification of comet tail moment. **p<0.01, ****p<0.0001, two-way ANOVA with Dunnett’s test for multiple comparisons. (I) Bar plot illustrating quantification of tail DNA percentage. ****p<0.0001, two-way ANOVA with Dunnett’s test for multiple comparisons.

**Figure S4.**
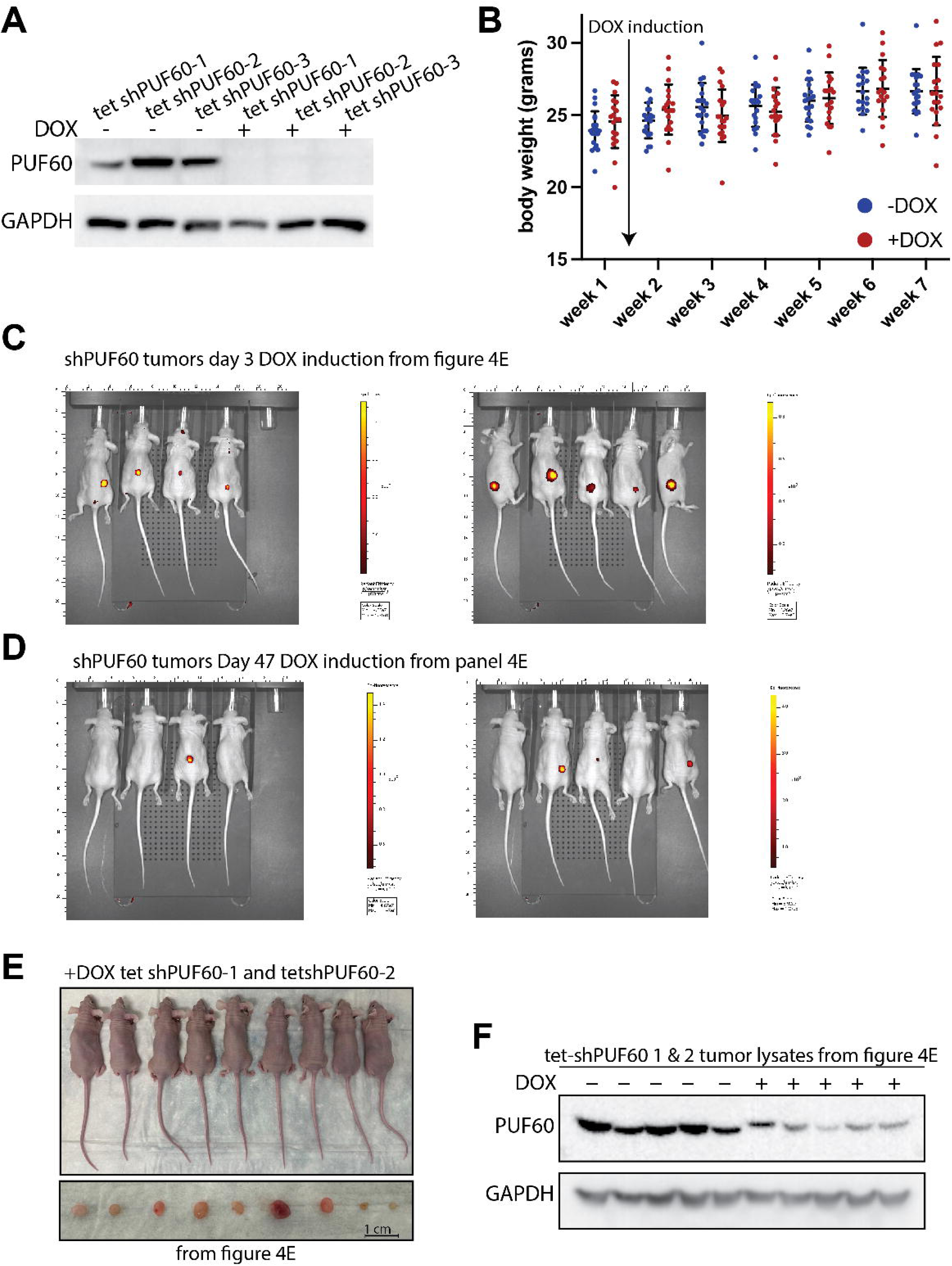
Depletion of PUF60 in TNBC cells significantly shrinks tumor volume, related to figure 4. (A) Western blot analysis of cell lysates from MDA-MB-231s transduced with 3 inducible shRNAs for PUF60 (tet-shPUF60-1, tet-shPUF60-2, tet-shPUF60-3). DOX treated cells are compared with untreated cells. (B) Body weights of female mice xenografted with tet-shPUF60-1 and tet-shPUF60-2. Measured twice weekly for 7 weeks. (C) Images of mice and excised tumors (Fig. 4E) 47 days following DOX induction compared with vehicle controls. (D) Western blot analysis of cell lysates from final tumors from (Fig. 4E) showing PUF60 protein expression in DOX-induced compared to vehicle control tumors. (E) In vivo imaging of mice bearing RFP-positive tumors on day 3 of DOX administration. (F) In vivo imaging of mice bearing RFP-positive tumors on day 47 of DOX administration.

**Figure S5.**
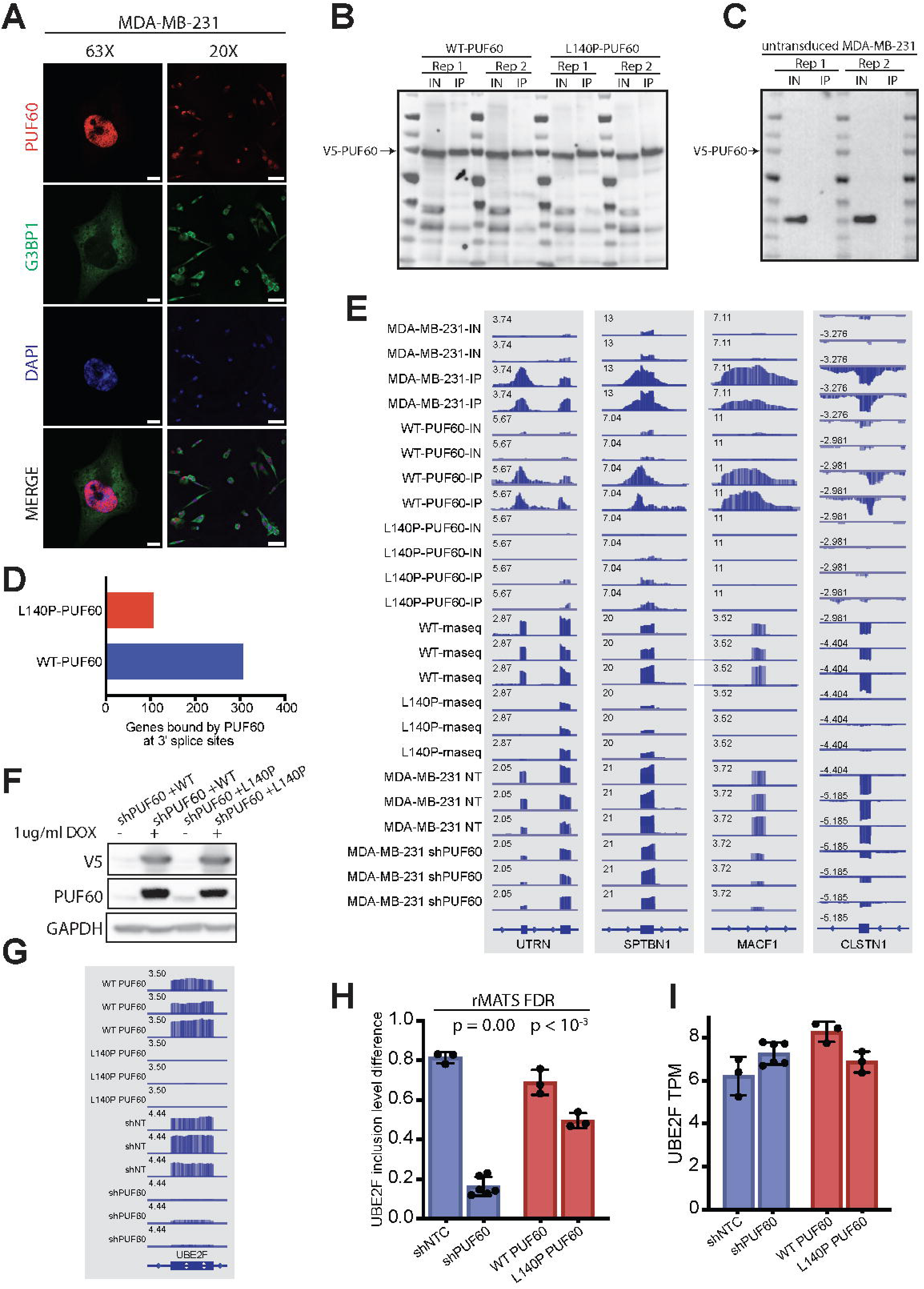
PUF60-activated exons are excluded following L140P substitution, related to figure 5. (A) Representative images showing PUF60 localization (red) in unperturbed MDA-MB-231s with cytoplasmic G3BP1 (green) and nuclear DAPI (blue) for reference. Images taken at 63x (Scale bar = 10 μm) and 20x (Scale bar = 100 μm) magnification. (B) Western blots of eCLIP immunoprecipitations for V5-tagged L140P-PUF60 and WT-PUF60 MDA-MB-231 cells. (C) Western blots of eCLIP immunoprecipitations for untransduced control MDA-MB-231 cells. (D) Barplot summarizing the number of genes bound at the 3’SS by WT-PUF60 or L140P-PUF60. (E) IGV browser tracks showing coverage of L140P-PUF60 and WT-PUF60 eCLIP signal relative to size-matched inputs. Tracks also show PUF60 knockdown and L140P-PUF60 RNA-seq signal relative to non-targeting shRNA and WT-PUF60 (respectively) at exons within cell cycle-associated transcripts. (F) Western blot analysis of cell lysates from MDA-MB-231s transduced with inducible, codon-optimized open reading frames (ORFs) encoding either L140P or wild-type (WT) PUF60 along with an inducible PUF60 shRNA. DOX treated cells are compared with untreated cells. (G) IGV browser tracks showing coverage of L140P-PUF60 RNA-seq signal relative to WT-PUF60 and exon 5 of UBE2F. (H) Bar plot summarizing UBE2F exon 5 inclusion level differences as determined by rMATS. False discovery rate (FDR) calculated by the rMATS log likelihood ratio. (I) Bar plots summarizing UBE2F TPMs.

**Figure S6.**
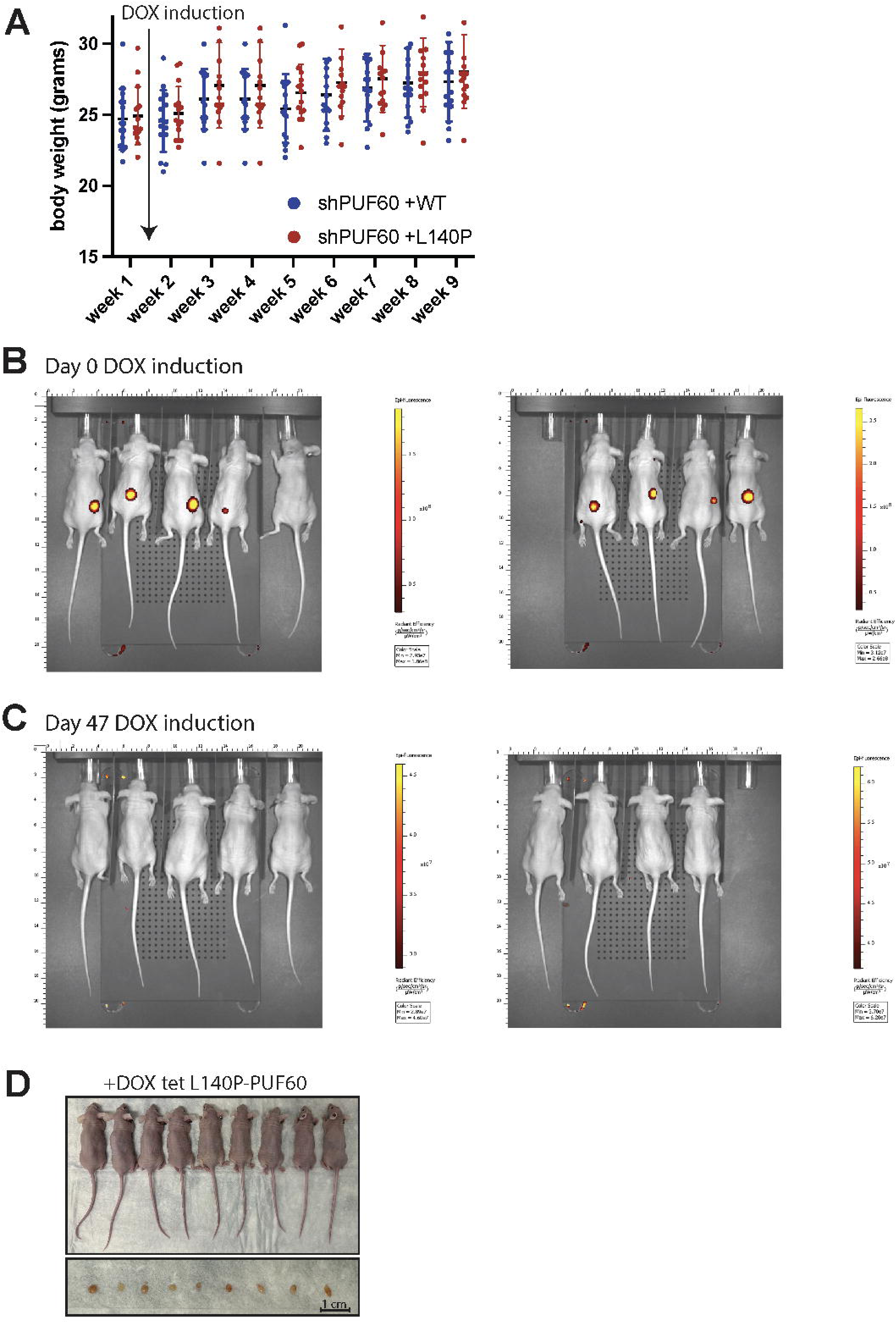
Silencing of PUF60 splice site activity shrinks tumor volume, related to figure 6. (A) Body weights of female mice xenografted with MDA-MB-231s stably expressing L140P-PUF60 and WT-PUF60. Measured twice weekly for 9 weeks. (B) In vivo imaging of mice bearing RFP-positive tumors on day 0 of DOX administration. (C) In vivo imaging of mice bearing RFP-positive tumors on day 47 of DOX administration. (D) Images of mice and excised tumors from (**Fig 6D**) 47 days following DOX induction.

